# Trustworthy agentic genomics through versioned skill libraries

**DOI:** 10.64898/2026.06.11.731523

**Authors:** Manuel Corpas, Alfredo Iacoangeli, Mathieu Bourdenx, Mahmoud Aldraimli, Nathan Skene, Segun Fatumo, Heinner Guio

## Abstract

Genomics is adopting autonomous AI agents that interpret genomes from natural-language instructions faster than it is building the means to trust them. We report the first large-scale controlled evaluation of where, in an agentic genomic pipeline, correctness must reside for the system to be trustworthy at clinical scale. Using pharmacogenomics, a domain where errors are measurable and sometimes lethal, we benchmarked nine frontier large language models across 44,550 scored evaluations on 110 pharmacogenomic cases, and tested model interpretation of real star-allele diplotypes from more than 7,000 individuals in three ancestrally diverse populations. Trustworthiness proved to be a property of pipeline architecture, not of the model. Letting the model reason was stochastic and unsafe, and grounding it in the correct guidelines by retrieval paradoxically increased lethal-class errors. Encoding the validated decision logic as a versioned skill and executing it as code made the pharmacogenomic mapping exact, auditable and identical across models, confining all residual error to a single input-interpretation step. On individual genomes, unguarded model interpretation degraded along an ancestry gradient; execution removes this gradient from the clinical mapping, relocating it to the auditable completeness of the input caller. This establishes a generalisable, auditable architecture for trustworthy agentic genome interpretation at scale.

**Highlights:**

- Correctness must be executed, not reasoned or retrieved, to be trustworthy
- Retrieval raises phenotype accuracy yet increases lethal-class errors; skills do not
- Execution makes the clinical mapping exact and model-invariant; error stays at input
- A deterministic input caller is the predicted route to all-correct emitted answers

**In brief:** Corpas and colleagues show that trustworthy agentic genome interpretation comes not from making language models reason correctly about biology, but from confining them to interpreting input while versioned, validated skills do the reasoning as executed code. Across nine large language models and 110 pharmacogenomics cases, executing the skill makes the clinical mapping deterministic, auditable and model-invariant.

**Significance:** Genomics is adopting autonomous, language-model-mediated agents faster than it is building the standards needed to trust them. On a pharmacogenomic benchmark with lethal-class consequences, we show that an agent’s trustworthiness is not a property of the model but of how the agent is constrained: correctness must be moved out of the stochastic model into a versioned skill executed as code, with the model confined to interpreting heterogeneous input. This gives the field a transferable architecture for trustworthy agentic genome interpretation, a predicted route to deploying it so that every emitted answer is correct (execute the validated skill, call the input deterministically, and abstain on the irreducible residual), and a way to develop genomic skills as validated, executable, versioned units rather than prompts. Following a validation framework described elsewhere, we use clinical-grade to mean determinism, auditability, traceability to versioned components and population-invariant performance, all achieved under skill-constrained execution. We distinguish two senses of population performance: the executed clinical mapping is population-invariant by construction, verified across European, Latin American and East African origin individuals, whereas the model’s interpretation of real, ancestrally diverse diplotypes is not, degrading along an ancestry gradient, which is precisely why the mapping must be executed rather than reasoned. We do not claim full clinical validation, which would additionally require non-canonical inputs, real-world genomic and clinical data, human comparators and multi-site concordance.

## Introduction

Genomics has entered a phase in which AI agents can autonomously discover, configure, execute and chain bioinformatics operations from natural-language instructions (1). This paradigm, agentic genomics, shifts the bottleneck in computational biology from building pipelines to validating them (1), and with it comes a new class of risk: agents can generate results faster than researchers can check them, and the most dangerous errors are the ones that look plausible but are wrong. A companion Perspective to this paper (1) sets out a tiered validation framework for agentic genomics and identifies the clinical-grade tier as the most demanding, requiring outputs that are deterministic, auditable, traceable to specific versioned components, and population-invariant. Here we operationalise that framework on a single clinical-task substrate, pharmacogenomic interpretation, to ask the question that decides whether agentic genome interpretation can be automated at scale: where in an agentic pipeline must correctness reside for the system to be trustworthy?

Two answers dominate practice. The first is to let the model reason from its training prior. The second, retrieval-augmented generation, grounds the model in retrieved evidence and lets it reason over that (2, 3, 4); large language models encode substantial clinical knowledge (5), and the reflex in biology is to surface the relevant guidelines and let the model apply them. This paper tests a third, structural answer: encode the validated decision logic in a versioned skill. In the agentic genomics framework, a skill is a self-contained, versioned unit of bioinformatics functionality whose declarative specification (SKILL.md) defines the analytical logic, expected inputs and outputs, and domain-expert decisions (1, 6). A skill can be used in two ways: the model can read its rules and reason to an answer, or the validated logic can be executed as code with the model confined to supplying input. That distinction turns out to be decisive.

Pharmacogenomics is a rigorous test domain: the genotype-to-phenotype-to-recommendation mapping is well-defined by the Clinical Pharmacogenetics Implementation Consortium (CPIC) Level A guideline corpus (7), errors are clinically measurable, and the task demands precise adherence to published guidance. The benchmark includes high-stakes drug-gene pairs with documented lethal consequences: DPYD variants where standard fluoropyrimidine dosing causes severe or fatal toxicity in approximately 0.1 to 0.5% of carriers (8); TPMT and NUDT15 deficiencies where standard thiopurine dosing causes lethal myelosuppression (9); CYP2C19 poor metabolisers where standard clopidogrel dosing fails to prevent stent thrombosis (10); and HLA risk alleles where exposure to specific drugs causes Stevens-Johnson syndrome, toxic epidermal necrolysis or hypersensitivity reaction (11, 12).

Concurrent work applies agents to genomic tasks but does not address this question. Recent systems generate pharmacogenomic recommendations from the literature through an agentic pipeline (13), or prioritise rare-disease variants for diagnosis at scale (14); both build end-to-end pipelines and report task accuracy. Neither isolates where in the pipeline correctness must reside, neither separates a model that reasons over a skill from one that executes it as code, and neither establishes determinism, auditability and model-invariance as properties of execution rather than of the model. That controlled dissociation, not the construction of another agentic system, is our contribution.

We benchmark nine frontier large language models along a five-step constraint gradient that progressively relocates correctness from the model into an executed skill: (i) free-prompted; (ii) retrieval-augmented from the CPIC corpus; (iii) skill-reasoning, in which the model applies a versioned skill’s rules and generates the answer; (iv) skill-execution, in which the model supplies a structured input and the validated skill computes the answer in code; and (v) an answer-supplied positive control. The gradient shows that trustworthiness is a property of where correctness sits, not of the model: the unconstrained and retrieval-grounded conditions are stochastic and unsafe; structured skills make capable models reliable and near model-invariant; execution makes the clinical mapping exact and confines all residual error to one auditable step; and a validated deterministic caller for that step closes the pipeline toward complete correctness among emitted in-scope answers. We frame the contribution as an architecture and an evaluation for trustworthy agentic genome interpretation at scale, not as a claim that models become clinically correct in isolation. Figure 1 summarises the study design.

**Figure 1.**
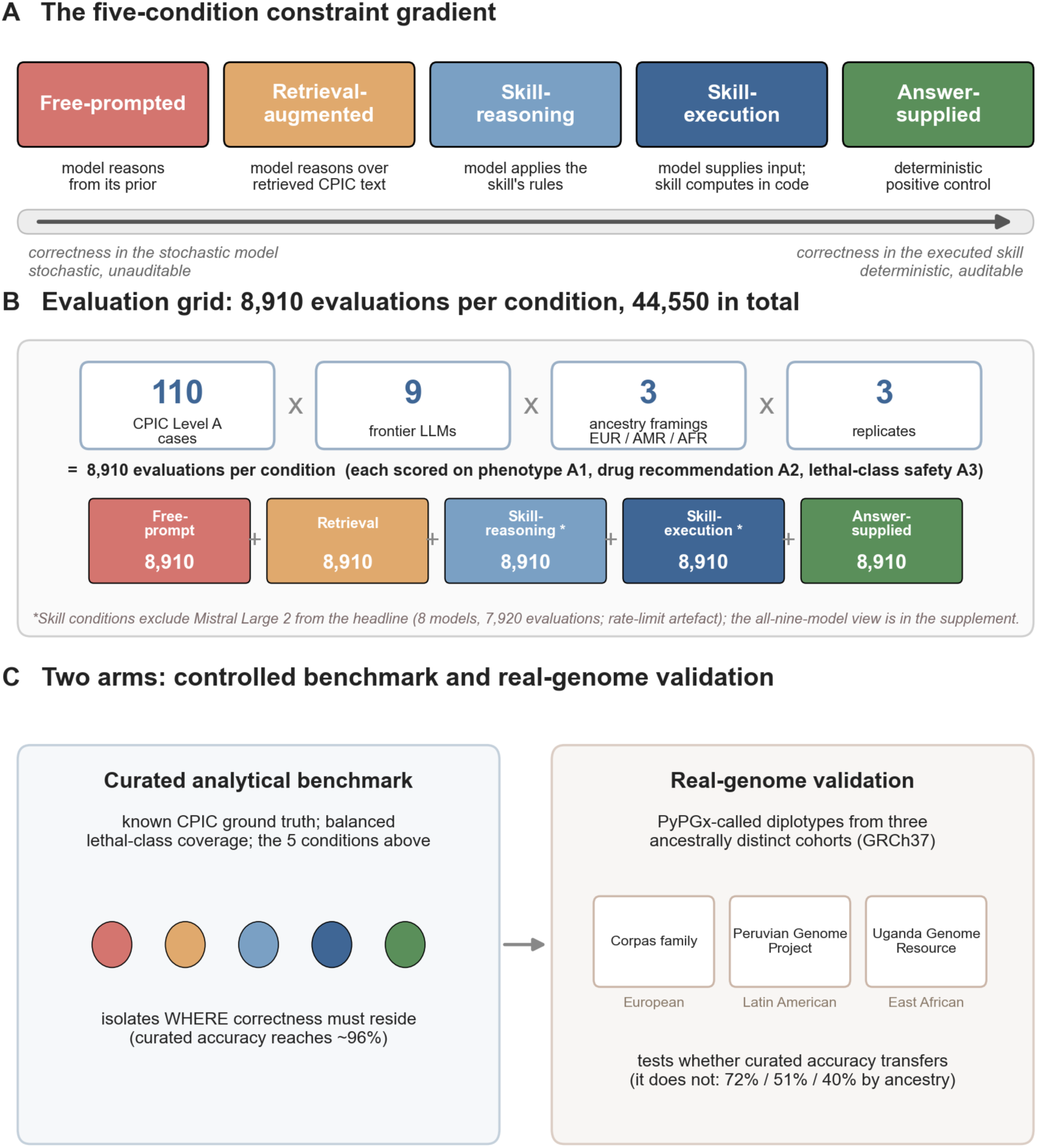
Study design. (A) The five-condition constraint gradient, ordered by how much correctness sits in the stochastic model versus in the executed skill: free-prompted, retrieval-augmented, skill-reasoning, skill-execution, and an answer-supplied positive control. (B) Every condition is evaluated on the identical grid: 110 CPIC Level A cases × nine frontier large language models × three ancestry framings (European, Latin American, East African) × three replicates (8,910 evaluations per condition, 44,550 in total), each scored on phenotype identification (A1), drug recommendation (A2) and lethal-class safety action (A3). The two skill conditions exclude Mistral Large 2 from the headline (rate-limit artefact); the all-nine-model view is reported in the supplement. (C) Two arms. A curated analytical benchmark with known CPIC ground truth (the five conditions) isolates where correctness must reside; a real-genome validation arm interprets PyPGx-called diplotypes from three ancestrally distinct cohorts (Corpas family, Peruvian Genome Project, Uganda Genome Resource) to test whether curated accuracy transfers.

## Results

### A constraint-gradient benchmark of agentic pharmacogenomic interpretation

We benchmarked nine frontier large language models (Claude Opus 4, Claude Sonnet 4, GPT-5.2, GPT-4.1, o3, o4-mini, Gemini 2.5 Flash, DeepSeek V3, Mistral Large 2) on 110 CPIC Level A cases spanning 21 markers and 35 gene-drug pairs. The three core conditions (free-prompted, retrieval-augmented, answer-supplied) were evaluated across three population contexts and three replicates, producing 26,730 evaluations. The two skill conditions (skill-reasoning, skill-execution) were run on all nine models across the same three-population, three-replicate grid, producing a further 17,820 evaluations; Mistral Large 2 is excluded from the reported skill-arm aggregate as a rate-limit artefact (it returned usable output on only 3.7% of skill-arm cells; Methods), so the headline skill figures are over eight models, and the all-nine-model aggregate is reported in the supplement. On the common set of eight models for which all five conditions yield valid output (Mistral fails only under the skill prompts), the constraint gradient is unchanged: free-prompted 79.6%, retrieval-augmented 89.1%, skill-reasoning 95.5%, skill-execution 93.3% and the answer-supplied control 100%. Outputs were scored on phenotype identification (A1), drug-specific recommendation (A2) and lethal-class safety action (A3) by a rigorous rescorer with a justified clinical-equivalence layer (Methods).

The five conditions form a gradient of constraint, ordered by how much correctness sits in the model versus in the executed skill (Figure 2).

**Figure 2.**
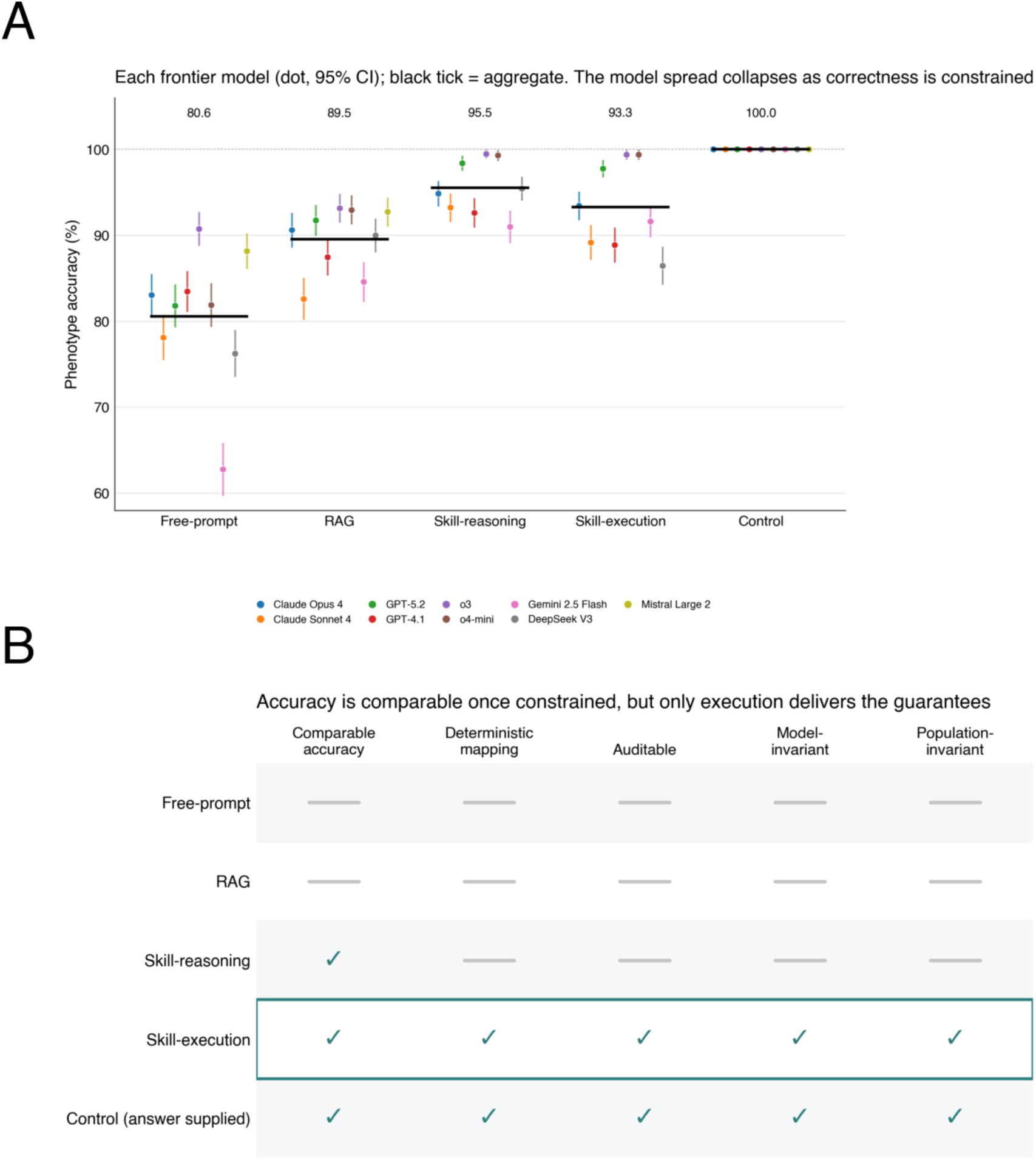
The constraint gradient locates trust. (A) Phenotype accuracy for each frontier model (coloured dot, with 95% confidence interval) across the five conditions, the black tick marking the aggregate (nine models for the free-prompted, retrieval-augmented and answer-supplied control conditions; eight models for the skill conditions, Mistral excluded). The per-model spread collapses as correctness is constrained; skill-reasoning (95.5%) and skill-execution (93.3%) reach comparable aggregate accuracy, and the gain over free-prompting (80.6%) comes from constraint rather than from execution per se. (B) Properties matrix: accuracy is comparable across skill-reasoning, skill-execution and the control, but only execution and the control deliver the four clinical-grade guarantees (deterministic mapping, auditability, model-invariance, population-invariance). Skill-execution (boxed) is the recommended deployment point.

### Unconstrained and retrieval-grounded agents are stochastic and unsafe

Free-prompted models reached 80.6% phenotype accuracy (95% CI 79.8 to 81.4) and committed 270 lethal-class safety errors on 1,096 parsed lethal-class cells, with three-of-three replicate consistency of only 82.7% and a 27.9-percentage-point spread between the weakest and strongest model (62.8% Gemini 2.5 Flash to 90.7% o3). The model was, in effect, the system, and an unreliable one.

Retrieval augmentation did not fix this. Phenotype accuracy rose to 89.5% (95% CI 88.8 to 90.1), but the lethal-class error rate rose from 24.6% (270 of 1,096 parsed lethal-class cells) to 36.6% (414 of 1,130) (two-proportion z = 6.1, P < 0.001) (Figure 3), drug-recommendation accuracy fell from 61.6% to 53.0%, and consistency remained imperfect at 93.8%. This safety regression does not depend on the scoring layer: the lethal-class error counts (270, 414) are identical under the baseline and clinical-equivalence scorers (Table S2). Providing the model with the correct guideline text raised one metric while degrading safety, the central anomaly we characterise next.

**Figure 3.**
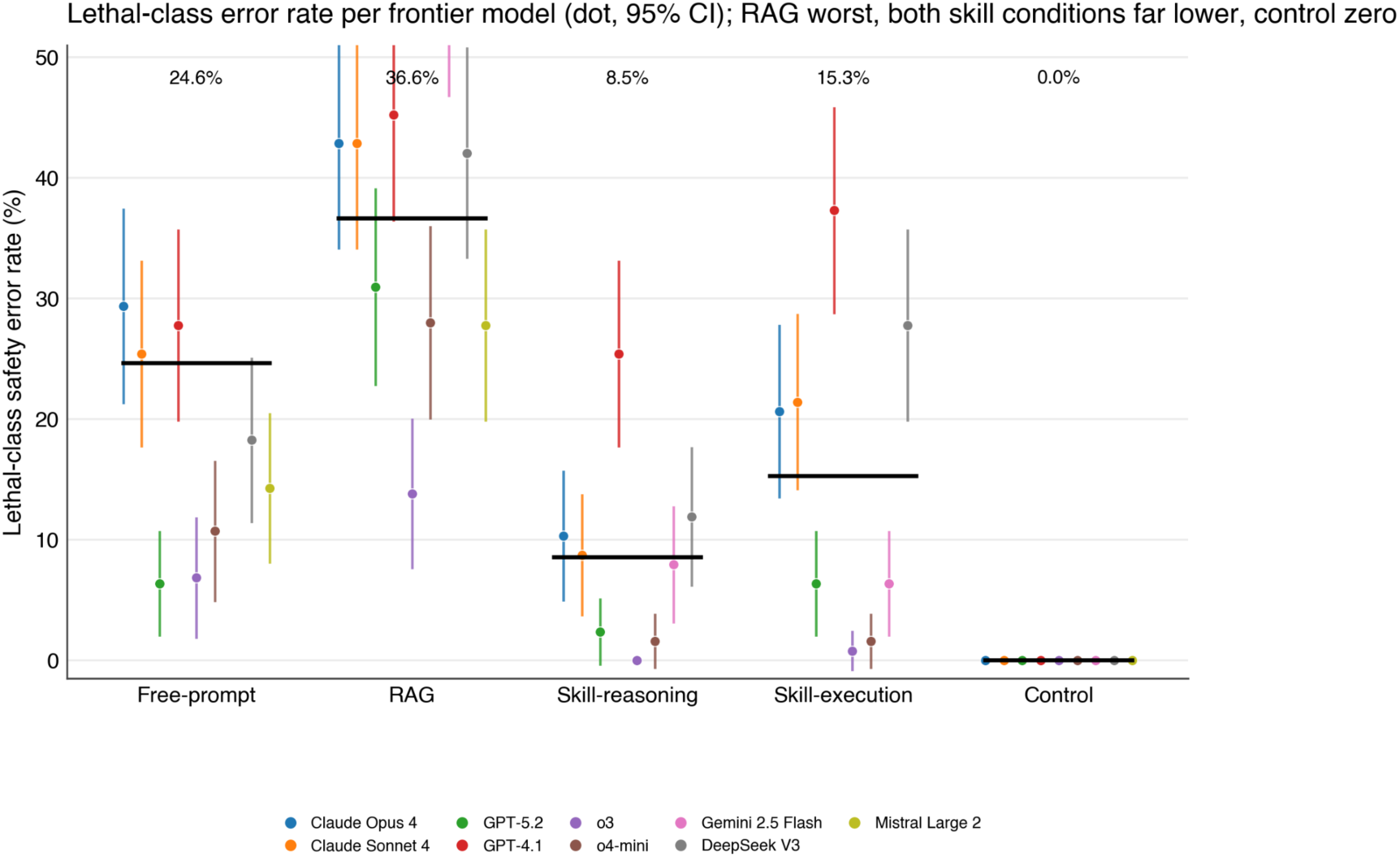
Retrieval raises phenotype accuracy yet increases lethal-class errors. Lethal-class safety error rate for each frontier model (coloured dot, with 95% confidence interval) across the gradient, the black tick marking the aggregate (nine models for the three-arm conditions, eight for the skill conditions). Retrieval augmentation is the worst condition for lethal-class errors; both skill conditions are far lower, with skill-execution’s residual modestly higher than skill-reasoning (15.3% versus 8.5%) but localised and removable by a deterministic caller (see text); only the deterministic answer-supplied control reaches zero. Surfacing the correct guideline text does not produce the correct safety action.

### Retrieval failure modes are architectural, not knowledge gaps

Two failure modes, absent from the other conditions, account for the retrieval regression. First, drug substitution: because CPIC guidelines are keyed on gene rather than gene-drug pair, a chunk retrieved for one drug carries recommendations for several, and models echo the wrong one. Drug substitution accounted for 470 of 973 drug-recommendation regressions (48.3%); 922 of 973 (94.8%) traced to chunk-structural confusion rather than clinical disagreement. The failure is a property of the index, not of retrieval as such: re-running the six worst-affected genes with finer (gene, drug)-keyed chunks eliminated drug substitution entirely, on 0 of 378 cells (Figure S2; Table S4). Second, information-without-action: on HLA risk-allele loci the model echoes the correct phenotype but softens the categorical AVOID recommendation, identifying HLA-B*57:01 status perfectly yet issuing AVOID on only a minority of lethal-class cells (Figure S3). A third pattern, correctness-by-coincidence, shows that 19.0% of free-prompted lethal-class correct-action cells reached the right recommendation through an incorrect phenotype; retrieval reduced this by about 40%, to 11.5%; only execution eliminated it (Figure S4). Notably, both characterised failure modes occur on loci other than TPMT and NUDT15, the two genes with documented corpus content gaps (Table S1), so the regression is a property of retrieval indexing rather than of those gaps. Pure outcome metrics conceal all three.

### Structured skills make capable models reliable and near model-invariant

When the validated decision logic was provided as a structured skill and the model applied it (skill-reasoning), phenotype accuracy rose to 95.5% (95% CI 95.1 to 96.0) with three-of-three replicate reproducibility of 96.4% (eight-model, three-population, three-replicate grid, Mistral excluded as a rate-limit artefact; Methods). The reasoning models reached the ceiling: o3 and o4-mini attained 99.3 to 99.5% with near-perfect reproducibility. Critically, the 27.9-percentage-point model spread of the free-prompted condition collapsed to single digits (Figure 4): a structured skill scaffolds weaker models up toward the strongest, so that model choice ceases to be the dominant determinant of accuracy. Skill-reasoning, however, is not deterministic by construction: the clinical answer still passes through the model’s generation, and a model’s stated reasoning is not a faithful guarantee of the computation behind it (15), so it is reliable rather than guaranteed.

**Figure 4.**
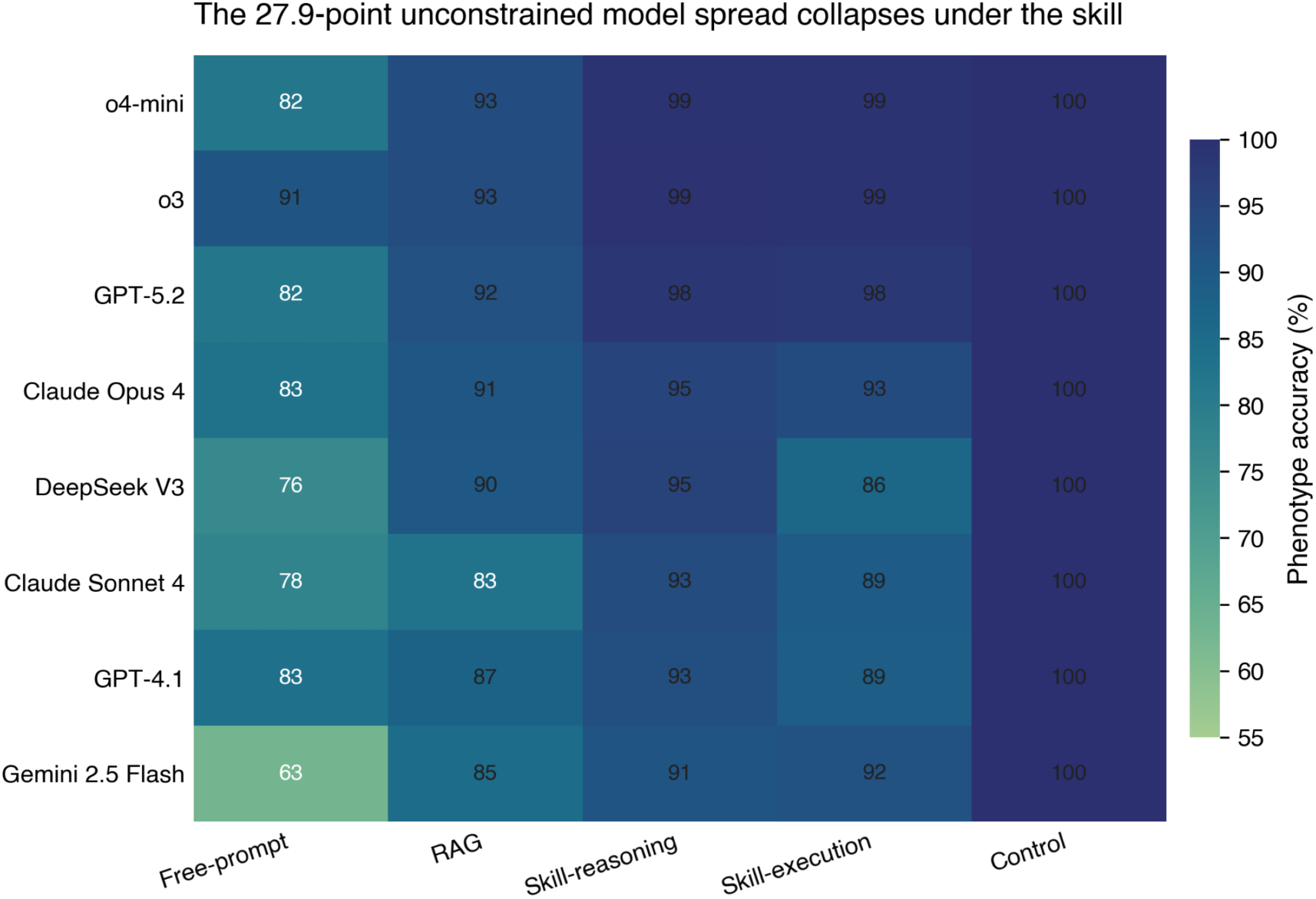
Skill execution collapses model differences. Per-model phenotype accuracy across the constraint gradient (eight models). The 27.9-percentage-point free-prompted spread between the weakest and strongest model (Gemini 2.5 Flash 62.8% to o3 90.7%) collapses to single digits once a structured skill supplies the decision logic, so model choice ceases to be the dominant determinant of accuracy and the cheapest compliant model reaches the ceiling.

### Execution makes the clinical mapping exact and localises all residual error to input interpretation

When the validated logic was executed as code, with the model confined to supplying a structured input (skill-execution), the behaviour changed in kind. Executing the skill on the correct diplotype produced the canonical phenotype, recommendation and safety action on every one of the 110 cases (A1 = A2 = A3 = 100%, zero lethal-class errors in this mapping-verification on the correct diplotype): the mapping is exact by construction. End-to-end phenotype accuracy was 93.3% (95% CI 92.7 to 93.8; three-population, three-replicate grid), and this number tracks, and is bounded below by, the accuracy of the model’s input-interpretation step: whenever the model called the correct diplotype, which it did on 92.0% of cells, the executed mapping was correct 100% of the time (Figure 5); on the remaining 8.0% of cells, where the call was wrong, the executed phenotype was nonetheless correct by coincidence in 15.8% of cases, so end-to-end accuracy (93.3%) marginally exceeds the correct-call rate (92.0%). A controlled-vocabulary input, in which the model selects a diplotype from the skill’s declared list, removed notation ambiguity (99.4% of outputs were in-vocabulary) and left a genuine input-interpretation residual of approximately 8%, concentrated in genes where star-allele calling from raw variants is hard (SLCO1B1, G6PD, VKORC1). The clinical mapping contributed no error.

**Figure 5.**
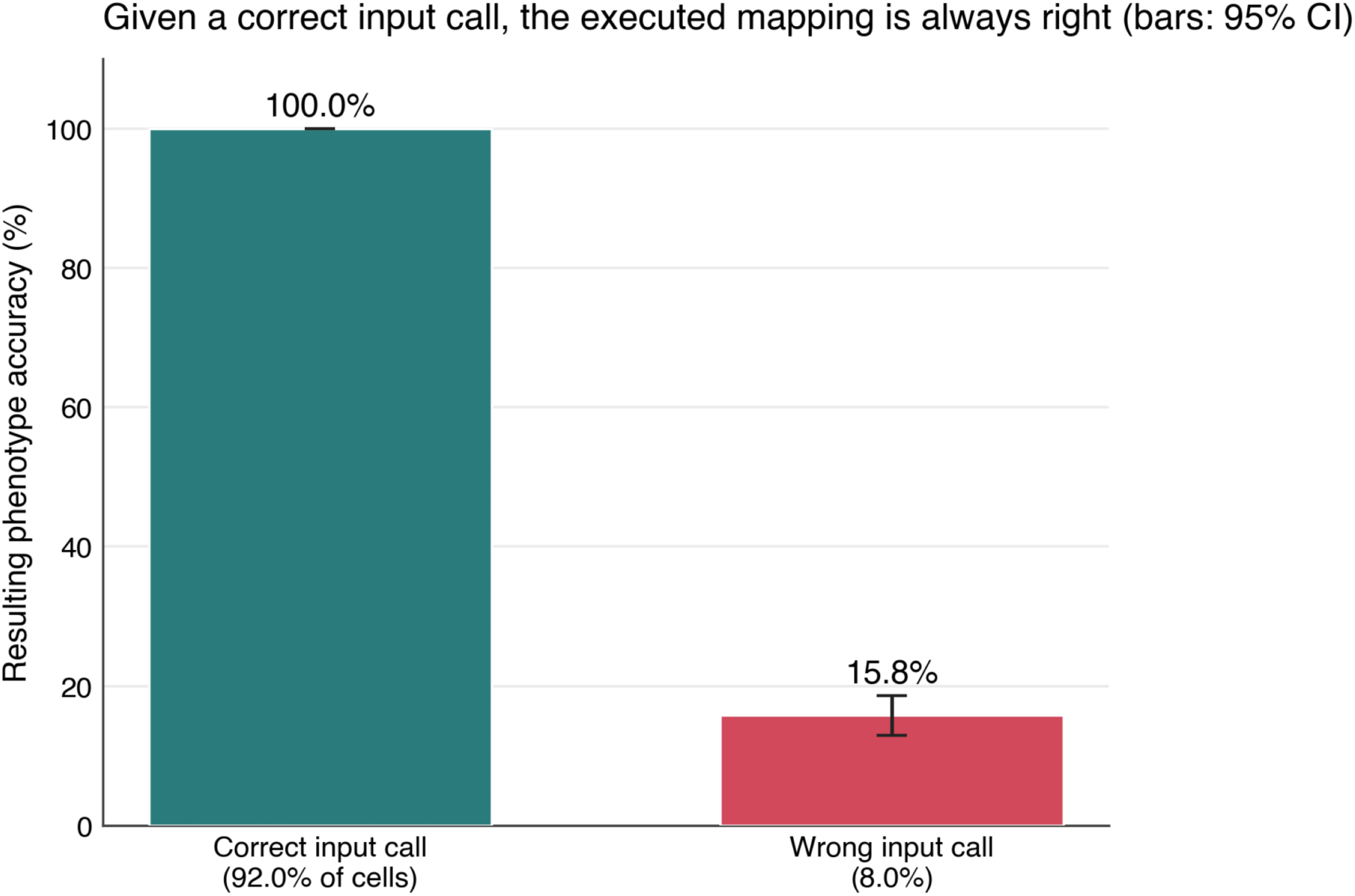
All residual error is the input call; the executed mapping is exact. Under skill-execution, end-to-end phenotype accuracy (93.3%) tracks input-interpretation accuracy (92.0%) and is bounded below by it. When the model called the correct diplotype, which it did on 92.0% of cells, the executed mapping returned the canonical phenotype 100% of the time; on the 8.0% of cells with an incorrect call the phenotype was correct only by coincidence (15.8%). The clinical mapping itself contributed no error. Error bars, 95% confidence intervals.

Skill-reasoning and skill-execution receive identical information, the same validated skill, and differ only in whether the model reasons over the rules or executes them as code, so the contrast isolates the locus of correctness rather than the amount of information supplied. Skill-reasoning was marginally more accurate than skill-execution (95.5% versus 93.3%) because, with the full rule table in context, a reasoning model can self-correct a loose input call, whereas execution commits to a single call and runs it rigidly. This is the central trade and the central point: execution does not win on raw accuracy, it wins on what clinical-grade requires, namely that the clinical mapping is provably deterministic and auditable and that the residual risk is localised to one measurable step rather than diffused across the model’s reasoning.

Skill-execution carries a higher lethal-class error rate than skill-reasoning (15.3% versus 8.5%), and the reason is the same trade that makes execution preferable. A reasoning model holding the full rule table can silently correct a loose input call, so its lethal-class residual is lower but unguaranteed and diffused across the model’s generation; execution commits to the single called diplotype and runs it rigidly, so a wrong input call propagates to a wrong, occasionally lethal, executed output. The residual is therefore higher under execution but localised to one measurable step, auditable, and removable by construction: a validated deterministic caller eliminates it at source and the abstain path withholds the rest. Reasoning’s lower rate buys no such guarantee. We recommend execution not because it minimises the raw lethal-class rate but because it is the only condition whose residual can be bounded, monitored and driven to zero.

### Toward 100% correctness among emitted answers: a deterministic input caller

Because the executed mapping is exact and the entire residual is the input call, the route to a fully correct pipeline is to make the input call deterministic. Substituting a validated diplotype caller, which maps raw variants to a diplotype with no generative step, removes the only error source by construction; the agent abstains on no-calls and ambiguous diplotypes. The real-genome arm here used PyPGx (0.26.0) as its reference caller; PharmCAT (16) is an example of a deployment-grade caller suited to this role. The achievable target is therefore not universal near-perfect clinical performance but a bounded guarantee: 100% correctness among emitted in-scope answers, with explicit abstention on no-calls, ambiguous alleles and any case outside the validated skill set. We do not claim near-100% accuracy on arbitrary real-world genomes; the real-genome arm below shows why such a claim would be false for model-mediated interpretation. The model’s role contracts to what it does well and uniquely at scale: parsing heterogeneous, natural-language input into the skill’s controlled vocabulary.

### Correctness resides in the executed contract, not the model

To establish that the skill conditions reflect contract execution rather than coincident model knowledge, we corrupted the skill in both directions. Across five lethal-class cases, three models and three replicates, with the phenotype and recommendation fields deliberately set to clinically wrong values, 90 of 90 responses executed the corrupted contract (88 echoed verbatim, two hedged while still echoing the unsafe fields) and none reverted to the canonical CPIC answer, in both the lethal-to-safe and safe-to-dangerous directions (Figure 6; Table S3). The answer-supplied control reached 100% identically across all nine models, vendors and generations (Table S2). Correctness is a property of the executed skill, and the model executes whichever contract it is given.

**Figure 6.**
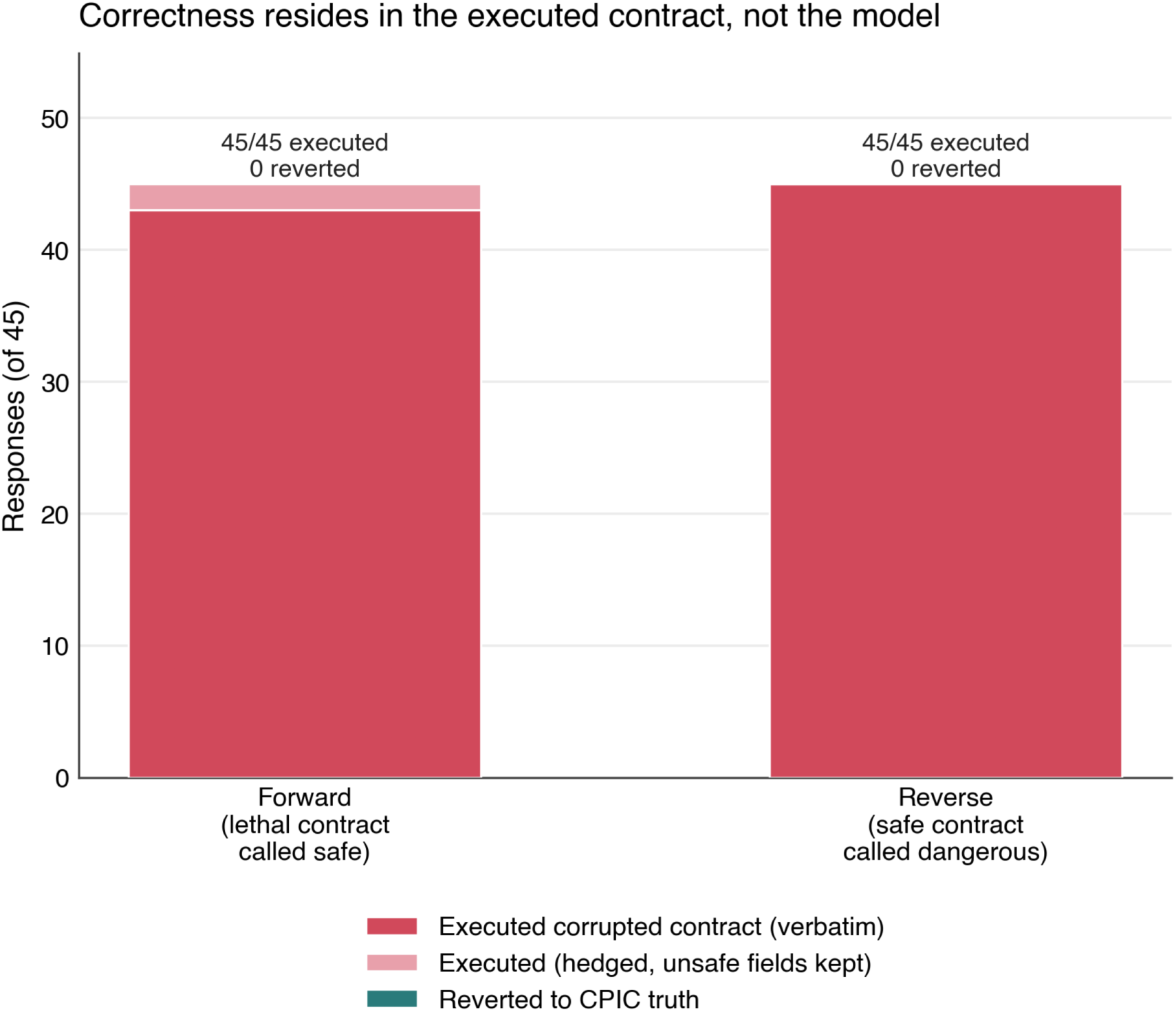
Correctness resides in the executed contract, not the model. Bidirectional adversarial scrambled-specification test across five lethal-class gene-drug pairs, three models and three replicates per direction (45 responses each). With the skill’s phenotype and recommendation fields deliberately corrupted, the model executed the corrupted contract in 90 of 90 responses (88 verbatim, two hedged while still emitting the unsafe fields) and never reverted to the canonical CPIC answer (0 of 90; the Wilson 95% upper bound on reversion is 4.1%), in both the lethal-to-safe and safe-to-dangerous directions.

### Population context

We evaluated all five conditions across three population contexts, European (Corpasome) (19), Latin American (Peruvian Genome Project) (20) and East African (Uganda Genome Resource) (21), using identical curated genotypes under population-specific clinical framing. Under this curated framing, accuracy was population-invariant, and the residual gap narrowed as correctness moved into the executed skill. This invariance is a property of the executed clinical mapping; it does not imply that the model can interpret real, ancestrally diverse diplotypes invariantly, which the real-genome arm below shows it cannot. The free-prompted EUR-AMR-AFR spread was 0.7 percentage points; under retrieval it was 0.9. For the skill conditions, measured on the eight non-Mistral models across all three populations and three replicates, the spread was 1.4 points for skill-reasoning (96.1%, 95.9%, 94.7%) and 1.7 points for skill-execution (93.8%, 93.9%, 92.2%), and the answer-supplied control was identical across populations by construction (Figure 7). These per-population values are pooled over the same three-replicate grid that anchors the headline accuracies (skill-reasoning 95.5%, skill-execution 93.3%). The small aggregate spread masks real, heterogeneous per-locus population gaps, shown in Figure S1. One honest caveat remains at the safety layer: lethal-class errors were higher for the East African context (skill-execution 56 versus 48 for European; skill-reasoning 41 versus 24), reflecting a slightly larger input-interpretation residual on the population whose star alleles (for example CYP2D6 *17) are least represented in reference panels. Population-invariance of the executed clinical mapping is therefore a property of execution rather than a tuning achievement, while the input-interpretation step retains the small, measurable, population-sensitive residual that the deterministic caller is designed to remove.

**Figure 7.**
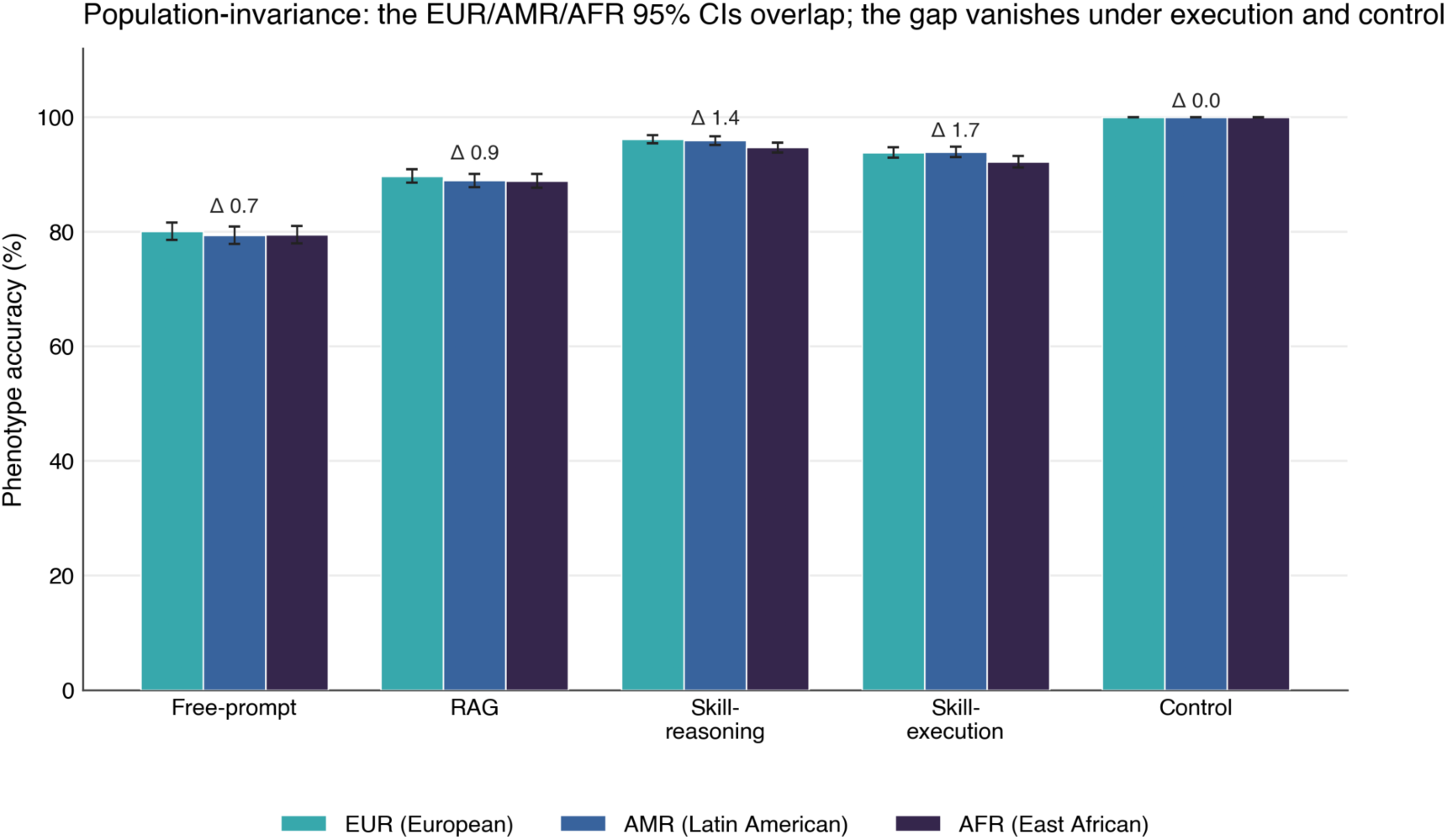
Population-invariance across three ancestry contexts. Phenotype accuracy for European (Corpasome), Latin American (Peruvian Genome Project) and East African (Uganda Genome Resource) framings of identical genotypes (eight models for the skill conditions). The EUR-AMR-AFR spread is 0.7 points free-prompted and remains under two points across all skill conditions (1.4 for reasoning, 1.7 for execution), and is zero for the answer-supplied control by construction. Error bars, 95% confidence intervals.

### Validation on real genomes across three populations

To test whether these results hold on real genomes rather than curated cases, we applied the agent to star-allele diplotypes actually observed in three ancestrally distinct cohorts, each called by a single deterministic caller (PyPGx) on GRCh37 so the comparison is fair: a European family (the Corpas family, five members) (19), the Peruvian Genome Project (an admixed Latin American cohort, 736 individuals across seven subpopulations) (20), and the Uganda Genome Resource (an East African cohort, 6,407 individuals) (21). This experiment deliberately tests the unsafe alternative, model-mediated interpretation of real diplotypes, to quantify why execution is required: it inverts the recommended architecture, in which the executed skill, not the model, performs the diplotype-to-phenotype mapping, precisely to measure the cost of letting the model interpret. The deterministic caller performed the input step (variants to diplotype) and the model was then made to perform the interpretation step (diplotype to CPIC phenotype), scored against the caller’s phenotype (Figure 8).

**Figure 8.**
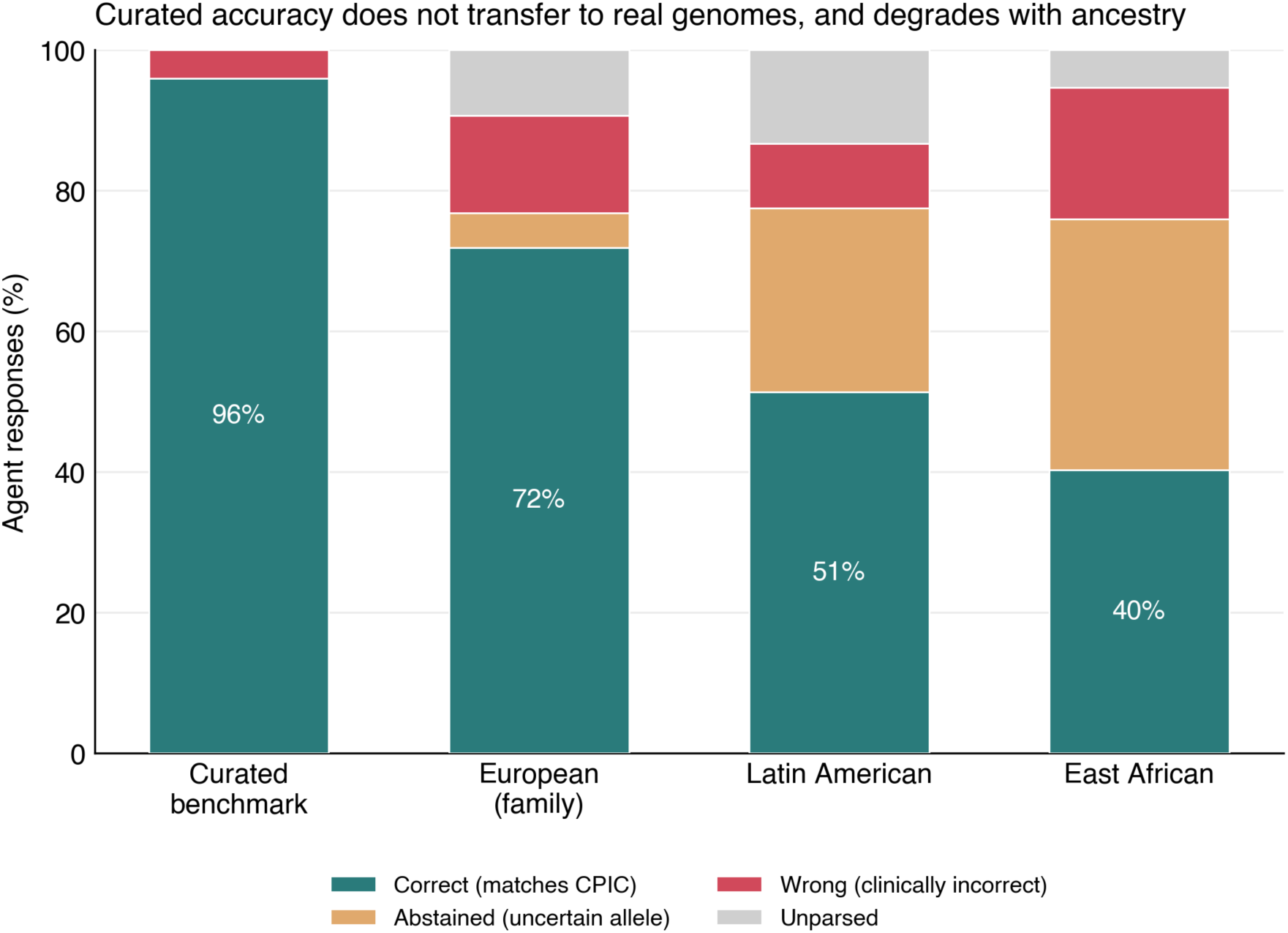
Curated accuracy does not transfer to real genomes and degrades with ancestry. Agent interpretation of real star-allele diplotypes observed in three cohorts, each called by the same deterministic caller (PyPGx, GRCh37), scored against the caller’s CPIC phenotype: a European family (Corpas family, five members), the Peruvian Genome Project (Latin American, 736), and the Uganda Genome Resource (East African, 6,407). Stacked bars show the proportion of agent responses that were correct, abstained (returned indeterminate on an uncertain allele), clinically wrong, or unparsed, against the curated-benchmark reference (96%). Correct interpretation falls along an ancestry gradient (72%, 51%, 40%) with abstention rising correspondingly (5%, 26%, 36%); the deterministic caller interpreted all diplotypes regardless of ancestry.

Curated accuracy did not transfer. Where the model reached 96% on the benchmark’s canonical diplotypes, on real diplotypes it correctly reproduced the phenotype for 72% of European (203 scored interpretations, from a single family of five related individuals and so not independent observations), 51% of Latin American (294) and 40% of East African (1,396); these denominators count distinct observed diplotype states interpreted once by each panel model, not individuals (Methods). The accuracy followed an ancestry-associated gradient, mirrored by abstention: the model declined to call (returning an indeterminate phenotype) on 5%, 26% and 36% of cases respectively, because real and especially African genomes carry star alleles under-represented in the literature the models were trained on (for example African-common CYP2C19 and DPYD alleles). The model was least able to interpret the genomes most distant from its European-centric training, and most likely to defer on them; clinically incorrect calls were also most frequent for the East African cohort (19% versus 14% European). Because the deterministic caller performed the input step here, it necessarily returned a diplotype for every case, so the relevant point is not that a caller never abstains but that the disparity we observe is a property of model interpretation rather than of the executed mapping. Substituting execution for model interpretation relocates the disparity upstream into the completeness of the caller’s allele-function tables, where African-common star alleles remain comparatively under-represented; that makes caller allele coverage a bounded, auditable engineering target rather than a diffuse model failure, though it does not by itself make the pipeline ancestry-fair. The cohorts differ in size, and the European anchor is a single family of five related individuals rather than an independent population sample, so we treat that point as illustrative and base the population comparison on the two large cohorts, where larger samples surface more rare alleles: Latin American 51% (n = 294, Wilson 95% CI 45.3 to 56.7) versus East African 40% (n = 1,396, Wilson 95% CI 37.4 to 42.6), a gap of 11 percentage points (two-proportion z = 3.5, P < 0.001), with the East African cohort showing both the most abstention and the most dangerous errors. The full pipeline is openly reproducible (Methods).

## Discussion

The five benchmarked conditions (free-prompted, retrieval-augmented, skill-reasoning, skill-execution and an answer-supplied control) are not points on a single accuracy gradient; they fail and succeed in qualitatively distinct ways, and ordered by how much correctness sits in the model versus in the executed skill they tell one story. As correctness moves out of the stochastic model and into the executed skill, accuracy rises, model differences collapse, lethal-class errors fall, and, decisively, the residual risk stops being diffuse and becomes localised, measurable and auditable. The conceptual lesson is not that large language models are intrinsically unsafe for clinical genomics, but that clinical-grade agentic implementations require correctness to reside in versioned skills, not in the model. We organise the implications around three questions: which principles generalise across genomics and biology, how to deploy a validated skill trustworthily at scale, and how to develop such skills.

### What principles generalise across genomics and biology

First, the trustworthy pattern is interpreter-plus-executed-skill, not reasoner. Scale wants the model, because it absorbs the heterogeneity of real inputs that breaks rigid pipelines; trust forbids the model from being the source of scientific truth, because anything it generates is stochastic and unauditable. The resolution is a division of labour: the model is the flexible natural-language interface and orchestrator, and validated, versioned, executable skills are the deterministic scientific core. This is what good bioinformatics already is; the contribution is locating the model’s safe place within it.

Second, the trust boundary is always the input-interpretation handoff. Our model-difference data make this concrete: the 27.9-point unconstrained spread collapsed under the skill and, in the executed condition, vanished from the clinical mapping and survived only at input interpretation (DeepSeek, for example, reasoned through the skill at 96.4% but called the diplotype at 84.5%). Wherever agents are deployed in biology, the fragile, model-dependent step is the conversion of messy reality into a structured call (entity disambiguation, variant notation, units, ontology mapping, tool selection), and that is where validation and monitoring belong.

Third, knowledge-in-context is necessary but not sufficient and can be actively harmful. Retrieval, the field’s default clinical-grounding strategy, raised one metric while increasing lethal-class errors, through architectural failure modes (drug substitution, information-without-action) rather than knowledge gaps. Surfacing the correct evidence does not produce the correct action; the decision logic must be encoded and executed, not inferred from retrieved text. Crucially, this also rules out the simplest alternative reading of the gradient, that it merely tracks how much information each condition is given: the condition with the most information in context, retrieval, is the least safe, while the two conditions given identical information, skill-reasoning and skill-execution, differ not in accuracy but in whether their correctness is guaranteed.

Fourth, reproducibility and auditability must be engineered, not expected. Science requires same-input-same-output and traceable provenance, which a generative model cannot provide; the reproducible, auditable core must be executed code, with the model confined to roles that are verified or non-load-bearing. Fifth, model-agnosticism is achievable and valuable: under execution the clinical output was identical across vendors and generations, so a validated skill can be revalidated once and run by any compliant model, decoupling clinical safety from model churn, including the silent behavioural drift documented for deployed models over time (17). Sixth, skills become the unit of trustworthy accumulation: progress accrues as a growing library of validated, executable, composable, independently auditable skills rather than as ever-larger models trained to know more biology.

Finally, the honest limit is a spectrum of codifiability. Pharmacogenomics is unusually tractable: discrete genotype, categorical phenotype, discrete action, a short guideline-codified mapping. The more a biological task reduces to a validatable executable contract, the more trustworthy an agent can be on it; the less it does, the more the agent’s outputs are hypotheses requiring human or experimental validation, with the skill providing scaffolding and auditability rather than guaranteed correctness. That spectrum tells the field where to deploy autonomous agents now (codifiable interpretation: pharmacogenomics, variant classification against fixed criteria, standardised QC, annotation) and where they remain assistive and exploratory (discovery, evidence weighing under uncertainty, hypothesis generation).

### How to deploy a validated skill at scale, I: the architecture for getting every emitted answer right

The results specify a deployment architecture for running the skill across many genomes agentically. Heterogeneous genomic input is normalised by the model into the skill’s input schema; a validated deterministic caller, not the model, maps variants to a diplotype, emitting an explicit no-call when ambiguous; the validated skill executes the diplotype-to-phenotype-to-action mapping in code; deterministic validation gates cross-check the model’s parsed inputs against the caller before anything is emitted; and the agent abstains and escalates to a human on no-calls, novel variants, or any gene-drug pair outside the validated skill set. Because the mapping is model-invariant, the orchestration model can be the cheapest available, or two models can be run on the input step and required to agree as a free reliability gate. The honest definition of success is that 100% of emitted responses are correct, with controlled abstention on the irreducible residual; the residual is small, localised to input interpretation, and exactly where quality control should sit. One cannot reach 100% by making the model reason better, because the model’s input call tops out below 100% and varies by model; one reaches it by executing the mapping, calling the input deterministically, validating the handoff, and deferring the rest.

### How to deploy a validated skill at scale, II: the model as a controlled, monitored operator

What makes a pipeline agentic, and what makes it trustworthy at scale, are operational rather than cognitive. In the configuration that is reliable, the model does not perform the science; it acts as an operator at the edges of the system. It ingests heterogeneous real-world input (variant call files, free-text requests), normalises it into a skill’s controlled vocabulary, selects the appropriate versioned skill from the library, and routes the call to the deterministic caller and the executed mapping. The scientific computation is executed code; the agent’s contribution is to run that code correctly, at high throughput, under control, and to watch it. This is a deliberately smaller role for the model than the field’s reflex assumes, and that contraction is the result, not a limitation: the trustworthy agent is a constrained one.

Three operational properties make this trustworthy where unattended model reasoning is not. First, control: every parsed input is validated against the deterministic caller before anything is emitted; two inexpensive models can be required to agree on the input call as a free reliability gate; and the agent abstains and escalates to a human on no-calls, novel or uncertain-function alleles, and any gene-drug pair outside the validated skill set. Because the executed mapping is exact, the only judgement the agent must police is whether to emit or to defer. Second, monitoring: at the throughput of a population biobank no human can audit individual calls, so the operator instruments the process, tracking per-skill call volumes, abstention rates, the size and location of the input-interpretation residual, and, critically, performance stratified by ancestry, so that a population-specific degradation surfaces as a monitored signal rather than as downstream harm. Third, adjustment: because the executed core is model-invariant, the orchestration model is swappable and can be monitored for drift across versions without revalidating the science.

Our real-genome results make the necessity of this loop concrete. When the model performs the interpretation by reasoning, curated accuracy (96%) does not transfer to real genomes and degrades with ancestry: it correctly reproduced the phenotype for 72% of European, 51% of Latin American and 40% of East African real diplotypes, with abstention rising along the same gradient (5%, 26%, 36%) as the model met more alleles outside its training, and with clinically incorrect calls most frequent for the East African cohort. At population scale an unmonitored reasoning pipeline would emit these as a large absolute number of plausible, undetected and inequitably distributed clinical errors. These are precisely the errors a controlled, monitored agent gates: the deterministic caller and the abstain path prevent the wrong call from being emitted, and ancestry-stratified monitoring flags the residual before it reaches patients. High throughput therefore does not relax the requirement for this architecture, it makes it mandatory, because control and monitoring are the only mechanisms whose reliability is a property of the system rather than of the model, and therefore the only ones that hold as the agent is run across millions of diverse real genomes.

### How to develop agentic genomic skills

For developers, the results change the unit of work from a prompt to a validated, executable artefact. A skill should ship as four separable parts: a declarative specification with provenance (source guideline and version, author, commit); a deterministic core that encodes the validated mapping as code and is the thing executed; a constrained input layer with an explicit controlled vocabulary and a deterministic mapper from raw data to it; and a first-class abstain path. The model sits only at the edges, normalising input and rendering output, never producing the result. The development workflow follows: encode the authoritative source as structured rules rather than prose; write the deterministic core test-first and require it to pass exactly; build or wrap the deterministic input mapper, because that is where the residual lives; add the model interface last, with outputs constrained to the schema and cross-checked against the deterministic components; and engineer abstention and provenance explicitly, versioning and regression-testing the skill against guideline updates as one would clinical decision-support software. Two consequences follow: maintain two test suites, one for logic correctness (deterministic, must pass exactly, validated without any model) and one for input-interpretation robustness (measured and monitored, accepted below 100% and bounded by validation and abstention); and treat the model as a swappable, monitored component, since the executed core is model-invariant.

### Limitations

The benchmark uses 110 synthetic cases with genotypes specified as text rather than variant call format files; we evaluate the analytical layer only. The free-prompted and retrieval-augmented prompts specify gene and genotype but not the individual drug, so their drug-recommendation and safety scores are evaluated at gene level (a correct recommendation for any drug of the gene counts); the skill conditions name the drug and are scored drug-specifically. Phenotype accuracy, the headline metric, is drug-independent and so is unaffected by this asymmetry. A drug-match audit confirmed that under the unspecified prompt, models frequently default to the gene’s canonical drug (for example clopidogrel for CYP2C19), which gene-level scoring correctly credits. The skill conditions were evaluated on the same three-population, three-replicate grid as the three core conditions; the executed mapping is exact by construction, and the input-interpretation residual is estimated over this full grid. The skills tested are decision tables for a guideline-codified domain; composition across multi-step agentic chains, where input-interpretation residuals and abstentions propagate, is untested. Mistral Large 2 retained elevated non-response even under a paced, rate-limited protocol (around 18% of skill-arm cells returned empty, with lower accuracy where it did respond), so it is reported separately and excluded from the headline skill-arm aggregate rather than treated as a clean result; its three-arm data, collected without that failure mode, are retained. The nine models reflect the frontier at the time of testing (early 2026); because the executed clinical mapping is model-invariant by construction, the central conclusions are robust to subsequent model releases, and the unconstrained-arm figures should be read as a snapshot of contemporaneous frontier capability rather than a fixed ceiling. We do not claim full clinical validation, which would additionally require non-canonical inputs, real-world genomic and clinical data, human comparators and multi-site concordance.

### Outlook

The immediate extensions are to integrate a validated deterministic caller and demonstrate 100% correctness among emitted in-scope answers with controlled abstention on the residual; to scale the skill conditions to the full population and replicate grid; and to test skill composition and selection across multi-step workflows. The same architecture should extend to variant interpretation, where the mapping is codified in ClinGen and ACMG-AMP guidance, to polygenic risk interpretation, and to rare-disease triage, where the validation properties demonstrated here must hold at every step of a longer chain.

## STAR Methods

Key resources are listed in Table S1. Case design, the three core conditions (free-prompted; retrieval-augmented from a curated PharmGKB-derived CPIC corpus (18); answer-supplied), the nine models, the rigorous scorer with its clinical-equivalence layer, the bidirectional adversarial scrambled-specification experiment, the (gene, drug)-keyed chunking comparator, and the quantification follow the locked three-arm protocol implemented in the released analysis code (see Data and code availability).

The two skill conditions instantiate a single pharmacogenomics skill specification spanning the 21 markers and 35 gene-drug pairs, whose validated decision logic (diplotype to phenotype and phenotype to recommendation) was curated from CPIC Level A guideline tables. In skill-reasoning, the decision rules are placed in the prompt and the model is given the genotype and must apply the rules to generate the phenotype and recommendation. In skill-execution, the model is given the genotype and the skill’s controlled vocabulary of valid diplotypes and must output one diplotype verbatim; the validated skill then computes phenotype and recommendation in code. Both conditions were run across all nine models on 110 cases, three populations (European, Latin American and East African) and three replicates, the same grid as the three core conditions, using identical genotypes under population-specific clinical framing and a shuffled, rate-limited execution order to avoid confounding population with provider throttling. Mistral Large 2 returned usable output on only 3.7% of skill-arm cells and is excluded from the reported skill-arm aggregate as a documented rate-limit artefact, so the headline skill figures are over eight models; the all-nine-model aggregate is reported in the supplement. Input-interpretation accuracy is the fraction of cells on which the executed mapping, applied to the model’s called diplotype, returns the canonical answer. Mapping exactness was verified by executing the skill on the canonical diplotype for every case. For the real-genome arm, star-allele diplotypes were called from each cohort with a single deterministic caller (PyPGx 0.26.0) on GRCh37 to keep the cross-population comparison fair: the Corpas family (publicly available 23andMe SNP-chip genotypes, lifted from NCBI36 (hg18) to GRCh37) (19), the Peruvian Genome Project (736 individuals, seven subpopulations) (20), and the Uganda Genome Resource (6,407 individuals, controlled access) (21). The model panel interpreted each observed diplotype to a CPIC phenotype, scored against the caller’s phenotype; uncertain-function diplotypes for which the caller returned an indeterminate phenotype were treated as abstention targets. The unit of analysis in this arm is the distinct (gene, diplotype) state observed in a cohort, not the individual: PyPGx calls are collapsed so that each distinct observed diplotype contributes once regardless of how many carriers share it, and each distinct state is then interpreted once by each of the eight panel models. The reported real-genome denominators (203 European, 294 Latin American, 1,396 East African) are these scored interpretations, that is distinct observed states multiplied by models, after removing states with an indeterminate reference phenotype and dropping connection errors. Larger and more diverse cohorts surface more distinct rare diplotypes, so this count grows with cohort diversity rather than linearly with sample size, and per-individual rates are therefore not comparable across cohorts; the five-member European family in particular contributes many shared haplotypes, so its 203 interpretations are not 203 independent observations. Analysis code, including the skill-condition runners, the mapping verification, and the full real-genome pipeline (real-genome-arm/: PLINK2 region extraction, PyPGx calling, agent panel and scoring, with pinned software versions and per-cohort data-access instructions), is available at https://github.com/manuelcorpas/24-AGENTIC-PGX-BENCHMARK (commit a6a8e9d). No individual-level genotypes are redistributed; the benchmark dataset is archived on Zenodo (DOI 10.5281/zenodo.20567742). Models and sampling. Frontier here denotes the leading models available at the time of testing, queried at fixed API snapshots: claude-opus-4-20250514, claude-sonnet-4-20250514, gpt-5.2, gpt-4.1, o3, o4-mini, gemini-2.5-flash and deepseek-chat, with mistral-large reported separately; the exact identifiers are listed in Table S1. No temperature or top-p override was applied: models ran at each provider’s default sampling, and the reasoning models (o3, o4-mini, GPT-5.2) do not expose a temperature parameter. Stochasticity was therefore measured rather than suppressed, through three-of-three replicate consistency. Output-token caps were set generously per experiment, well above the length of a phenotype-and-recommendation answer and not binding on answer content: 500 to 600 tokens for the three-arm conditions (free-prompted, retrieval-augmented, answer-supplied) and the adversarial test, 400 for the skill conditions, and 120 for the single-line real-genome interpretation step; reasoning models that reject a chat-endpoint token cap used a 1,500 to 2,000-token completion budget, and Gemini 2.5 Flash used 1,000 to 4,096 (raised after an observed truncation). Requests were spaced per provider to respect rate limits; exact per-experiment values are in Table S1.

### Statistical reporting

Headline accuracies are point estimates over the stated denominators (110 cases per model and condition for the skill arm, pooled across replicates and, where indicated, across models), reported with binomial Wilson 95% confidence intervals: free-prompted 80.6% (79.8 to 81.4, n = 8,738), retrieval-augmented 89.5% (88.8 to 90.1, n = 8,790), skill-execution 93.3% (92.7 to 93.8, n = 7,920) and skill-reasoning 95.5% (95.1 to 96.0, n = 7,920). The free-prompted-to-retrieval increase in lethal-class errors (24.6% to 36.6%) and the Latin-American-to-East-African real-genome accuracy gap (51% to 40%) are each tested with a two-proportion z-test (z = 6.1 and z = 3.5 respectively, both P < 0.001). The per-condition and per-model counts needed to recompute exact intervals are released with the data. Remaining differences are interpreted qualitatively, by kind of failure and by whether a guarantee holds by construction, because the executed-mapping comparisons are deterministic by design.

### Ethics and data governance

The real-genome cohorts were analysed under their respective ethics approvals and consent frameworks. The Corpas family 23andMe genotypes are openly published with participant consent. The Peruvian Genome Project data were collected under the project’s ethics approvals and were used here with the project’s permission. The Uganda Genome Resource is controlled-access data governed by a data transfer and access agreement and was analysed solely under its terms. No individual-level genotypes from any cohort are redistributed, and only aggregate, non-identifying results are reported.

### Data and code availability

This study uses three human cohorts, obtained through their respective access routes. No individual-level genotypes are redistributed in this paper or in the associated repository.

### European (Corpas family)

The 23andMe SNP-chip genotypes of the five family members (19) are publicly available on figshare (“23andMe SNP chip genotype data”, https://doi.org/10.6084/m9.figshare.92682) and are openly downloadable without restriction. The conversion from NCBI36 (hg18) to GRCh37 used in this study is reproduced by the pipeline.

### Latin American (Peruvian Genome Project)

The cohort is described in the Peruvian Genome Project paper (20), with per-subpopulation diplotype-to-phenotype and per-individual recommendation tables in that paper’s supplementary materials. Genotype data are available from the project (INBIOMEDIC and UTEC, H.G.) on reasonable request, subject to the project’s data-sharing terms.

### East African (Uganda Genome Resource)

The UGR (21) is controlled-access genomic data governed by a data transfer and access agreement and is not redistributable. Access is granted on application to the data custodians (MRC/UVRI and LSHTM Uganda Research Unit, and the Uganda Medical Informatics Centre). Approved users receive the GRCh37 genotype set that this pipeline takes as input.

### Code and derived data

All analysis code, including the skill-condition runners, the mapping-verification step, and the full three-population real-genome pipeline (region extraction, PyPGx calling, agent panel, scoring, and figure generation), with pinned software versions and per-cohort data-access instructions, is publicly available at https://github.com/manuelcorpas/24-AGENTIC-PGX-BENCHMARK (commit a6a8e9d). The curated benchmark dataset (no individual-level genotypes) is archived on Zenodo (https://doi.org/10.5281/zenodo.20567742); the raw, rescored three-arm and adversarial datasets are archived separately on Zenodo (https://doi.org/10.5281/zenodo.20567743). Aggregate per-population result tables that underlie the figures contain no individual-level data and are included in the repository.

## Author contributions

M.C.: Conceptualisation, Methodology, Software, Investigation, Formal analysis, Writing - original draft, Writing - review and editing, Supervision, Funding acquisition. A.I.: Methodology, Writing - review and editing. M.B.: Methodology, Writing - review and editing. M.A.: Writing - review and editing. N.S.: Writing - review and editing. S.F.: Methodology (population stratification), Writing - review and editing. H.G.: Methodology (Latin American cohort context), Writing - review and editing.

## Declaration of interests

M.C. is the founder of ClawBio, whose SKILL.md specification format this study evaluates, and the validation-tier framework on which the study draws is defined in the companion Perspective under review (reference 1). M.C. and H.G. are associated with GENEQ Global Ltd. The other authors declare no competing interests.

## Acknowledgements

We thank CPIC and the PharmGKB team for the publicly available guideline corpus, and the contributors to the ClawBio open-source skill library. No external funding was received for this work.

## Notes

https://github.com/manuelcorpas/24-AGENTIC-PGX-BENCHMARK

https://doi.org/10.5281/zenodo.20567742

https://doi.org/10.5281/zenodo.20567743

### Competing Interest Statement

Manuel Corpas and Heinner Guio are associated with GENEQ Global, a company developing genomics-based health intelligence tools. All other authors declare no competing interests.

## References

1. Corpas, M., Fatumo, S., and Guio, H. (2026). Agentic genomics: from pipeline automation to autonomous validation. Cell Genomics. Manuscript under review.

2. Lewis, P., Perez, E., Piktus, A., Petroni, F., Karpukhin, V., Goyal, N., Küttler, H., Lewis, M., Yih, W.-t., Rocktäschel, T., et al. (2020). Retrieval-augmented generation for knowledge-intensive NLP tasks. Adv. Neural Inf. Process. Syst. 33, 9459–9474. 10.48550/arXiv.2005.11401.

3. Xiong, G., Jin, Q., Lu, Z., and Zhang, A. (2024). Benchmarking retrieval-augmented generation for medicine. In Findings of the Association for Computational Linguistics: ACL 2024, pp. 6233–6251. 10.48550/arXiv.2402.13178.

4. Zakka, C., Shad, R., Chaurasia, A., Dalal, A.R., Kim, J.L., Moor, M., Fong, R., Phillips, C., Alexander, K., Ashley, E., et al. (2024). Almanac: retrieval-augmented language models for clinical medicine. NEJM AI 1, AIoa2300068. 10.1056/AIoa2300068.

5. Singhal, K., Azizi, S., Tu, T., et al. (2023). Large language models encode clinical knowledge. Nature 620, 172–180. 10.1038/s41586-023-06291-2.

6. Corpas, M. (2026). ClawBio: Bioinformatics-Native AI Agent Skill Library (v0.5.0). Zenodo. 10.5281/zenodo.19420648.

7. Relling, M.V., and Klein, T.E. (2011). CPIC: Clinical Pharmacogenetics Implementation Consortium of the Pharmacogenomics Research Network. Clin. Pharmacol. Ther. 89, 464–467. 10.1038/clpt.2010.279.

8. Henricks, L.M., Lunenburg, C.A.T.C., de Man, F.M., Meulendijks, D., Frederix, G.W.J., Kienhuis, E., Creemers, G.-J., Baars, A., Dezentjé, V.O., Imholz, A.L.T., et al. (2018). DPYD genotype-guided dose individualisation of fluoropyrimidine therapy in patients with cancer: a prospective safety analysis. Lancet Oncol. 19, 1459–1467. 10.1016/S1470-2045(18)30686-7.

9. Relling, M.V., Schwab, M., Whirl-Carrillo, M., et al. (2019). Clinical Pharmacogenetics Implementation Consortium guideline for thiopurine dosing based on TPMT and NUDT15 genotypes: 2018 update. Clin. Pharmacol. Ther. 105, 1095–1105. 10.1002/cpt.1304.

10. Lee, C.R., Luzum, J.A., Sangkuhl, K., et al. (2022). Clinical Pharmacogenetics Implementation Consortium guideline for CYP2C19 genotype and clopidogrel therapy: 2022 update. Clin. Pharmacol. Ther. 112, 959–967. 10.1002/cpt.2526.

11. Mallal, S., Phillips, E., Carosi, G., et al. (2008). HLA-B*5701 screening for hypersensitivity to abacavir. N. Engl. J. Med. 358, 568–579. 10.1056/NEJMoa0706135.

12. Chung, W.-H., Hung, S.-I., Hong, H.-S., Hsih, M.-S., Yang, L.-C., Ho, H.-C., Wu, J.-Y., and Chen, Y.-T. (2004). A marker for Stevens-Johnson syndrome. Nature 428, 486. 10.1038/428486a.

13. Zack, M., Savinkov, A., Stupichev, D., Moore, A., Sokolov, D., Trifonov, I., Yankovskiy, A., Reshetnikov, K., Ydyrysova, N., and Gobbs, A. (2026). An agentic AI system for automated pharmacogenomic recommendation generation. npj Digit. Med. 10.1038/s41746-026-02590-w.

14. Kara, M., Gungor, A.F., Kuday, S.E., Ozcelik, O., Ceylaner, S., Dereli, F., and Ozden, F. (2026). Deep agentic variant prioritisation for expert-level genetic diagnosis fast at scale. medRxiv. 10.64898/2026.02.17.26346421.

15. Turpin, M., Michael, J., Perez, E., and Bowman, S.R. (2023). Language models don’t always say what they think: unfaithful explanations in chain-of-thought prompting. In Advances in Neural Information Processing Systems 36 (NeurIPS 2023). 10.48550/arXiv.2305.04388.

16. Sangkuhl, K., Whirl-Carrillo, M., Whaley, R.M., Woon, M., Lavertu, A., Altman, R.B., Carter, L., Verma, A., Ritchie, M.D., and Klein, T.E. (2020). Pharmacogenomics Clinical Annotation Tool (PharmCAT). Clin. Pharmacol. Ther. 107, 203–210. 10.1002/cpt.1568.

17. Chen, L., Zaharia, M., and Zou, J. (2024). How is ChatGPT’s behavior changing over time? Harvard Data Sci. Rev. 6. 10.1162/99608f92.5317da47.

18. Whirl-Carrillo, M., Huddart, R., Gong, L., Sangkuhl, K., Thorn, C.F., Whaley, R., and Klein, T.E. (2021). An evidence-based framework for evaluating pharmacogenomics knowledge for personalized medicine. Clin. Pharmacol. Ther. 110, 563–572. 10.1002/cpt.2350.

19. Glusman, G., Cariaso, M., Jimenez, R., Swan, D., Greshake, B., Bhak, J., Logan, D.W., and Corpas, M. (2012). Low budget analysis of Direct-To-Consumer genomic testing familial data. F1000Research 1, 3. 10.12688/f1000research.1-3.v1.

20. Guio, H., Sanchez, C., Borda, V., Jaramillo-Valverde, L., Caceres, O., Padilla, C., Trujillo, O., Poterico, J.A., Silva-Carvalho, C., Horton, M., et al. (2025). The Peruvian Genome Project: expanding the global pool of genome diversity from South America. Front. Genet. 16, 1614021. 10.3389/fgene.2025.1614021.

21. Gurdasani, D., Carstensen, T., Fatumo, S., Chen, G., Franklin, C.S., Prado-Martinez, J., et al. (2019). Uganda Genome Resource enables insights into population history and genomic discovery in Africa. Cell 179, 984–1002. 10.1016/j.cell.2019.10.004.

