## Supplementary materials for "Trustworthy agentic genomics through versioned skill libraries"

### Supplementary Information

#### Condition nomenclature

This study evaluates a five-condition constraint gradient (main text Figure 2): free-prompted, retrieval-augmented, skill-reasoning, skill-execution, and an answer-supplied positive control. The supplementary tables and figures below characterise the three core conditions that share the locked three-arm protocol, namely free-prompted, retrieval-augmented and the answer-supplied control (8,910 evaluations per condition per population set). The condition labelled with_spec in earlier dataset releases is the answer-supplied control referred to here and in the main text. The two skill conditions, skill-reasoning (the model applies a versioned skill's rules) and skill-execution (the validated logic is executed as code), were run on the same eight-model, three-population, three-replicate grid as the three core conditions and are reported in the main text; they are not part of the three-arm tables below.

#### Prompt specifications for the five conditions

Every condition was given the same clinical task, mapping a patient's genotype to a CPIC phenotype, drug recommendation and safety action, across the same 110 cases. The conditions differ only in where the decision logic resides: in the model's training prior (free-prompted), in retrieved guideline text (retrieval-augmented), in a structured rule table the model applies (skill-reasoning), in executed code with the model supplying only the input (skill-execution), or supplied directly (answer-supplied positive control). The verbatim prompt templates are below; curly-brace fields are populated per case, and the population framing (cohort name and population context) is identical across conditions. The full instantiated prompts, the per-gene skill rule tables and the CPIC guideline excerpts are in the analysis repository.

##### 1. Free-prompted

```
A patient from the {cohort name} ({cohort description}) has the following pharmacogenomics
genotype:

Gene: {gene}
Genotype: {genotype}

Based on CPIC guidelines, provide:
1. Star allele diplotype
2. Metaboliser phenotype
3. Drug recommendation
4. Clinical hazard if mismanaged
5. Population-specific considerations

Format as exactly 5 lines:
DIPLOTYPE: [answer]
PHENOTYPE: [answer]
DRUG: [drug]: [recommendation]
HAZARD: [hazard or None]
POPULATION: [population-specific note]
```

##### 2. Retrieval-augmented

```
You are a pharmacogenomics interpretation system. Use the CPIC guideline excerpt below to
decide the diplotype, phenotype, drug recommendation, hazard, and population-specific note
for the patient.

## CPIC guideline excerpt (retrieved for gene: {gene})
```

```
{gene-level CPIC excerpt, including the full recommendation table}

## Patient

Gene: {gene}
Genotype: {genotype}
Patient cohort: {cohort name} ({cohort description})

## Output (exactly 5 lines, no preamble)

DIPLOTYPE: [your answer]
PHENOTYPE: [your answer]
DRUG: [drug]: [recommendation]
HAZARD: [clinical hazard]
POPULATION: [population-specific note]
```

##### 3. Skill-reasoning

```
You are executing a ClawBio pharmacogenomics skill. Determine the patient's diplotype from
the genotype, then APPLY the skill rules below EXACTLY to output the phenotype and drug
recommendation.

{validated skill rule table for the gene: diplotype -> phenotype, and (drug, diplotype) ->
recommendation}

## Patient
Cohort: {cohort name} ({cohort description})
Gene: {gene}
Genotype: {genotype}
Drug: {drug}
Population context: {population note}

## Output (4 lines only)
DIPLOTYPE: [called diplotype]
PHENOTYPE: [from Rule 1]
DRUG: [from Rule 2]
HAZARD: [clinical hazard]
```

##### 4. Skill-execution (the model supplies only the input; a validated skill then computes phenotype and recommendation in code)

```
You are the input-interpretation step of a ClawBio pharmacogenomics agent. Map the
patient's genotype to exactly ONE diplotype from the controlled list below, copying its
text VERBATIM. Do not invent notation. A downstream validated skill computes phenotype and
recommendation.

Patient cohort: {cohort name} ({cohort description})
Gene: {gene}
Valid diplotypes (choose one, copy verbatim):
{controlled vocabulary: the valid diplotypes for the gene}

Patient genotype: {genotype}
Population context: {population note}

## Output (1 line only)
DIPLOTYPE: [exact text of one list item]
```

##### 5. Answer-supplied positive control

```
You are executing a ClawBio pharmacogenomics skill. Follow the specification EXACTLY.

## SKILL.md Specification
Gene: {gene}
Input: {genotype}
Patient cohort: {cohort name} ({cohort description})
Diplotype: {correct diplotype}
Phenotype: {correct phenotype}
```

```
Activity score: {correct activity score}
Drug recommendation: {correct recommendation}
Population context: {population note}

## Output (5 lines only):
DIPLOTYPE: {correct diplotype}
PHENOTYPE: {correct phenotype}
DRUG: {correct recommendation}
HAZARD: [clinical hazard]
POPULATION: {population note}
```

As a worked example, for CYP2D6 with genotype rs3892097 T/T (a *4/*4 case) and drug codeine: the free-prompted condition receives only the gene and genotype; retrieval-augmented additionally receives the gene's CPIC excerpt, which contains the codeine recommendation among several drug rows; skill-reasoning additionally receives the validated CYP2D6 rule table, including *4/*4 -> Poor Metaboliser and (codeine, *4/*4) -> AVOID; skill-execution receives the controlled vocabulary of CYP2D6 diplotypes and must output exactly one, after which the validated skill computes the phenotype and recommendation in code; and the answer-supplied control receives the answer itself (*4/*4, Poor Metaboliser, AVOID codeine) to echo. The genotype and the clinical question are identical throughout; only the scaffolding differs, which is the variable the constraint gradient manipulates.

**Population framing (identical across all five conditions)**
The cohort name, description and per-case population context are inserted identically into every condition's prompt; only the population label and its note change. For the CYP2D6 and codeine example the three tested contexts are:

**European:** European family cohort (Corpasome project); European ancestry, whole-genome sequencing. Population context: CYP2D6*4 ~20% in Europeans; PM ~7%. *1xN (UM) ~1-3%.

**Latin American:** Peruvian Genome Project; admixed Latin American, 7 indigenous and mestizo subpopulations. Population context: CYP2D6*4 ~10% in admixed Latin Americans; *10 and *17 more common in indigenous populations.

**East African:** Uganda Genome Resource; East African, 6,407 whole-genome sequences. Population context: CYP2D6*4 ~6% in Africans; *17 (reduced function) ~20-35%, often missed by EUR-centric panels.
Full prompts, rule tables and CPIC excerpts: https://github.com/manuelcorpas/24-AGENTIC-PGX-BENCHMARK (commit a6a8e9d).

#### Table S1. Key resources table

| CATEGORY | | |
|---|---|---|
| **REAGENT or RESOURCE** | **SOURCE** | **IDENTIFIER** |
| **Software and algorithms** | | |
| Claude Opus 4 | Anthropic | Model: claude-opus-4-20250514 |

| | | |
|---|---|---|
| Claude Sonnet 4 | Anthropic | Model: claude-sonnet-4-20250514 |
| GPT-5.2 | OpenAI | Model: gpt-5.2 |
| GPT-4.1 | OpenAI | Model: gpt-4.1 |
| o3 | OpenAI | Model: o3 |
| o4-mini | OpenAI | Model: o4-mini |
| Gemini 2.5 Flash | Google | Model: gemini-2.5-flash-preview-04-17 |
| DeepSeek V3 | DeepSeek | Model: deepseek-chat |
| Mistral Large 2 | Mistral AI | Model: mistral-large-2407 |
| ClawBio skill library (v0.5.0) | This paper | github.com/ClawBio/ClawBio; Zenodo: https://doi.org/10.5281/zenodo.19420648 |
| Benchmark, rescoring and figure code | This paper | github.com/manuelcorpas/24-AGENTIC-PGX-BENCHMARK (commit a6a8e9d) |
| Python 3.11 | Python Software Foundation | https://www.python.org |
| **Deposited data** | | |
| CPIC guideline corpus (v2026.04) | CPIC | https://cpicpgx.org; accessed 2026-04-15 |
| PharmGKB clinical annotations bundle | PharmGKB | https://www.pharmgkb.org; release 2026-04-09 |
| Raw, rescored three-arm and adversarial datasets | This paper | Zenodo: https://doi.org/10.5281/zenodo.20567743 |

**Table S1. Key resources.** Each large language model was queried via its public API at the model version listed, during the benchmark window. Decoding used default temperature throughout (no temperature or top-p override). Output-token caps were set per experiment, well above answer length and non-binding: 500 to 600 for the three-arm conditions and the adversarial test, 400 for the skill conditions, and 120 for the real-genome interpretation step; reasoning models used max_completion_tokens of 1,500 to 2,000, and Gemini 2.5 Flash used maxOutputTokens of 1,000 to 4,096 (raised after an observed truncation). The CPIC corpus for the cpic_rag arm is PharmGKB-derived; 19 of 21 genes are fully verified, with documented content gaps for TPMT and NUDT15.

#### Table S2. Headline metrics under the baseline and clinical-equivalence scorers

| Metric | Baseline scorer (10-rescore-v3.py) | Clinical-equivalence scorer (10b-rescore-v3-clinical-equivalence.py) |
|---|---|---|
| Aggregate phenotype accuracy A1 (n = 8,738) | 79.8% | 80.6% |
| Aggregate drug-recommendation accuracy A2 | 61.6% | 61.6% |
| Aggregate safety-action accuracy A3 | 96.9% | 96.9% |

| | | |
|---|---|---|
| Lethal-class A1 (n = 1,096) | 67.3% | 72.4% |
| Lethal-class A3 errors (count) | 270 | 270 |
| Three-of-three replicate consistency | 82.9% | 82.7% |
| **cpic_rag (retrieval-augmented)** | | |
| Aggregate phenotype accuracy A1 (n = 8,790) | 82.9% | 89.5% |
| Aggregate drug-recommendation accuracy A2 | 53.0% | 53.0% |
| Aggregate safety-action accuracy A3 | 95.3% | 95.3% |
| Lethal-class A1 (n = 1,130) | 61.2% | 86.3% |
| Lethal-class A3 errors (count) | 414 | 414 |
| Three-of-three replicate consistency | 92.8% | 93.8% |
| **answer-supplied control** | | |
| Aggregate phenotype accuracy A1 (n = 8,910) | 100.0% | 100.0% |
| Aggregate drug-recommendation accuracy A2 | 100.0% | 100.0% |
| Aggregate safety-action accuracy A3 | 100.0% | 100.0% |
| Lethal-class A1 (n = 1,134) | 100.0% | 100.0% |
| Lethal-class A3 errors (count) | 0 | 0 |
| Three-of-three replicate consistency | 100.0% | 100.0% |

**Table S2. Headline metrics under the rigorous baseline scorer and the clinical-equivalence scorer.** The equivalence layer modifies phenotype identification (A1) only, promoting natural-language risk phrasing to the canonical CPIC tier label, gated on locus context, and can only raise scores. Drug-recommendation accuracy (A2), safety-action accuracy (A3) and the lethal-class A3 error counts (270, 414, 0) are identical under both scorers, so the safety conclusions do not depend on the equivalence layer. The layer is generous to the retrieval-augmented arm, raising its aggregate A1 by 6.6 and its lethal-class A1 by 25.1 percentage points; even so, that arm retains more lethal-class errors than the free-prompted baseline (414 versus 270) and lower A2 (53.0% versus 61.6%). The answer-supplied control's 100% ceiling and determinism are identical under both.

#### Table S3. Bidirectional adversarial scrambled-specification test

| Direction | Cases (n=5 each) | ECHO scrambled | HEDGE (still unsafe) | Override to CPIC truth |
|---|---|---|---|---|
| Forward (lethal -> safe) | DPYD, CYP2D6, HLA-B*57:01, TPMT, CYP2C19 | 43 / 45 | 2 / 45 | 0 / 45 |
| Reverse (safe -> dangerous) | DPYD, CYP2D6, HLA-B*57:01, TPMT, CYP2C19 | 45 / 45 | 0 / 45 | 0 / 45 |
| Combined | | 88 / 90 | 2 / 90 | 0 / 90 |

**Table S3. Contract-faithfulness is symmetric.** Forward: five lethal-class specifications corrupted to look safe (Poor Metaboliser to Normal Metaboliser or Positive to Negative; AVOID to standard dosing). Reverse: five genuinely safe specifications corrupted to look dangerous (Normal Metaboliser or Negative to the dangerous phenotype; standard dosing to AVOID). Each direction: three models (Claude Opus 4, GPT-5.2, DeepSeek V3), three replicates per case (45 responses). In both directions every response executed the corrupted contract and none reverted to the canonical CPIC recommendation. Scripts: 14-adversarial-scrambled-spec.py, 14b-adversarial-reverse-spec.py; data: v3_adversarial_scrambled.json, v3_adversarial_reverse.json.

#### Table S4. Drug substitution under gene-keyed versus (gene, drug)-keyed chunking

| Gene | Gene-keyed chunks (drug-substitution %) | (gene, drug)-keyed chunks (drug-substitution %) | n (new run) |
|---|---|---|---|
| CYP2D6 | 67.8% | 0.0% | 120 |
| CYP2C19 | 61.3% | 0.0% | 120 |
| CYP2C9 | 44.6% | 0.0% | 54 |
| UGT1A1 | 36.4% | 0.0% | 36 |
| SLCO1B1 | 37.0% | 0.0% | 36 |
| IFNL3 | 100.0% | 0.0% | 12 |
| **All six genes** | **57.5%** | **0.0%** | **378** |

**Table S4. Drug substitution is eliminated by finer chunking.** The retrieval-augmented arm was re-run for the six worst-affected genes with (gene, drug)-keyed chunks (the model receives only the CPIC annotation section for the queried drug) on three models (Claude Opus 4, GPT-5.2, DeepSeek V3), the European population and two replicates (378 calls). Drug substitution (queried drug name absent from the parsed DRUG field, the same definition used for the gene-keyed arm) fell from 57.5% to 0 of 378 cells, on every gene. The drug-substitution failure is therefore a property of gene-keyed retrieval indexing, not of retrieval augmentation in general. Gene-keyed percentages are computed on the European cells of the locked three-arm dataset. Script: 16b-rag-genedrug-chunking.py; data: v3_rag_genedrug_chunking.json.

### Supplementary Results

#### Population context

Population-stratified phenotype accuracy showed only a small aggregate EUR-AFR gap under free prompting (81.0%, 80.4%, 80.3% for EUR, AMR, AFR) that retrieval augmentation did not narrow, over substantial but non-monotonic per-locus heterogeneity (per-gene EUR-AFR differences spanning about -10 to +10 points; Figure S1). The answer-supplied control removed both by construction: every case received an identical output regardless of population annotation, giving 100.0% accuracy on each population and zero per-locus delta.

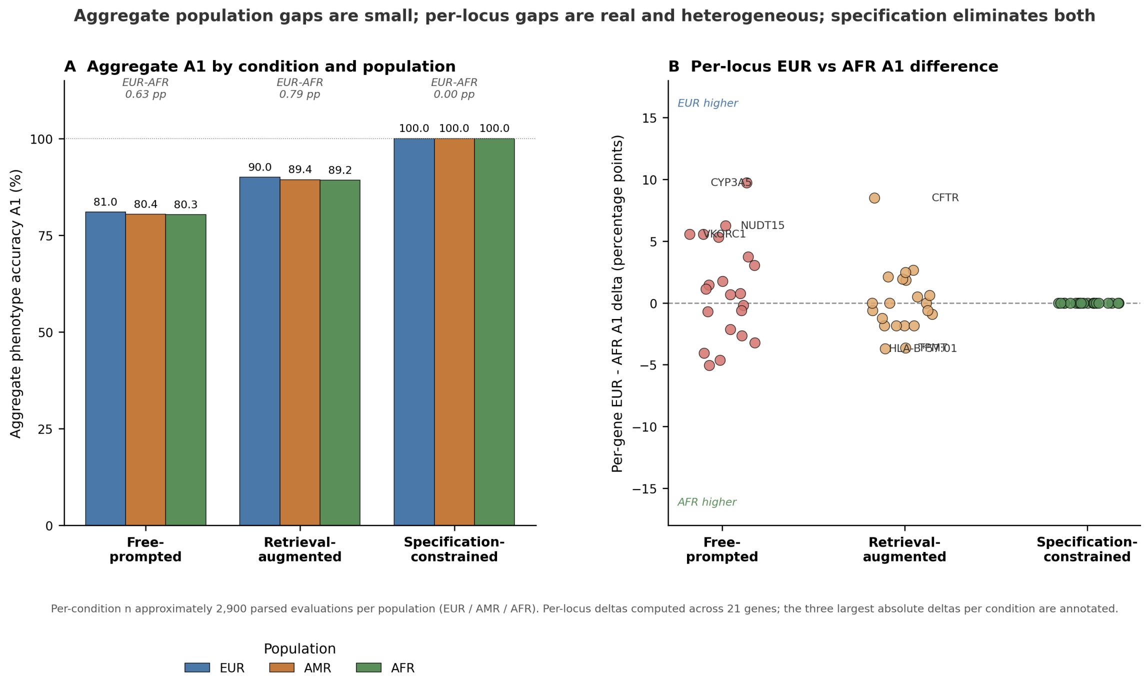


**Figure S1.** Aggregate population gaps are small; per-locus gaps are real and heterogeneous; the answer-supplied control eliminates both. (A) Aggregate phenotype accuracy (A1) by condition and population (EUR, AMR, AFR). EUR-AFR percentage-point gaps annotated above each cluster. (B) Per-gene EUR minus AFR A1 deltas by condition across 21 genes; the three largest absolute deltas per condition are annotated. The answer-supplied control produces 100.0% A1 on every population with zero per-locus delta on every gene.

Per-gene robustness

Across loci, the three conditions partition the 21 markers into deployment categories (Figures S1 and S3): loci where retrieval cleanly helps (for example HLA-B*58:01, CYP3A5); multi-drug loci where retrieval introduces drug substitution (CYP2D6, CYP2C19, CYP2C9, UGT1A1, SLCO1B1, IFNL3); and HLA risk-allele loci where retrieval introduces information-without-action (HLA-B*57:01, HLA-B*15:02, HLA-A*31:01). The answer-supplied control is uniformly safe across all three categories.

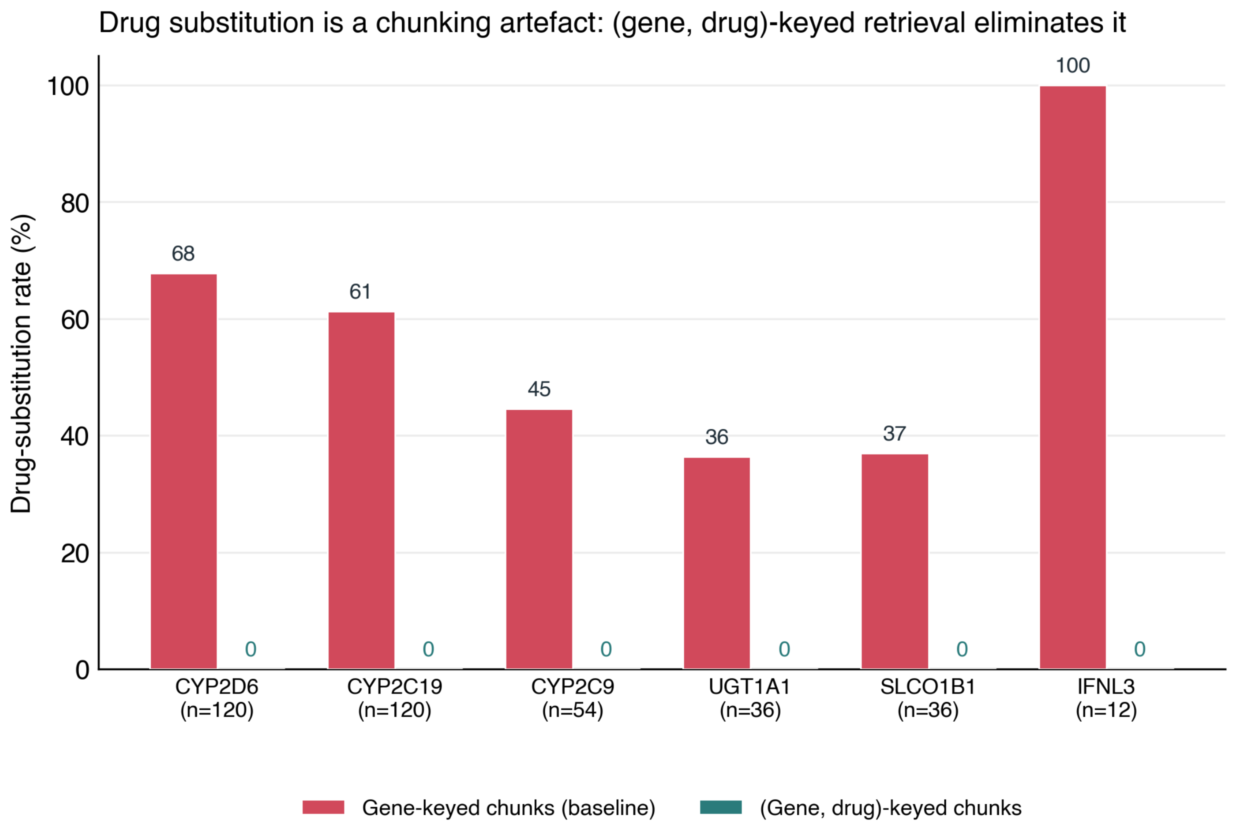


**Figure S2. Drug substitution is a chunking artefact.** Drug-substitution rate on the six worst-affected genes under the published gene-keyed chunks versus finer (gene, drug)-keyed chunks (retrieval-augmented condition, three models, European framing, two replicates; 378 calls). A gene-keyed chunk carries CPIC recommendations for several drugs at once and the model echoes the wrong one; restricting the chunk to the queried drug eliminates drug substitution entirely (0 of 378 cells).

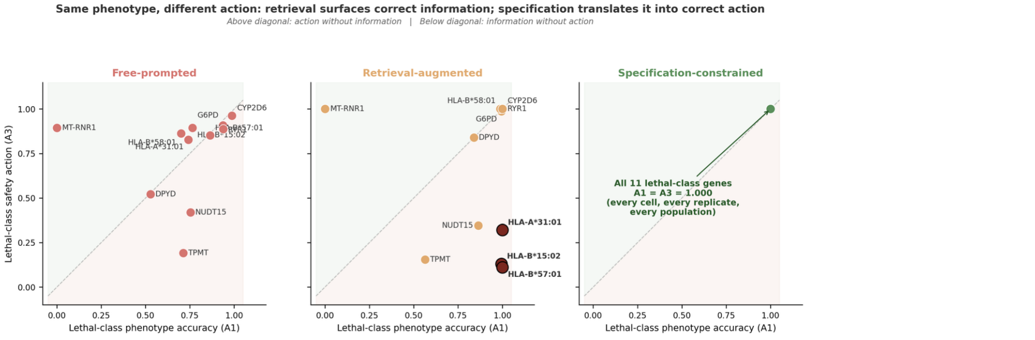


11 lethal-class genes (14 lethal cases). Per-gene means computed across all parsed lethal-class cells (populations and replicates pooled). Highlighted points in the Retrieval-augmented panel (HLA-B*57:01, HLA-B*15:02, HLA-A*31:01) show the canonical information-without-action pattern: phenotype correctly identified, AVOID action not produced.

**Figure S3.** Information without action. Lethal-class genes plotted at their mean phenotype-identification accuracy (A1) against mean clinical-safety action accuracy (A3), one panel per condition; the diagonal is alignment. Points below the diagonal identify the phenotype correctly but fail to issue the canonical AVOID action. Under retrieval, HLA-B*57:01, HLA-B*15:02 and HLA-A*31:01 show this information-without-action pattern; the answer-supplied control aligns A1 and A3 at (1, 1) on every lethal-class gene.

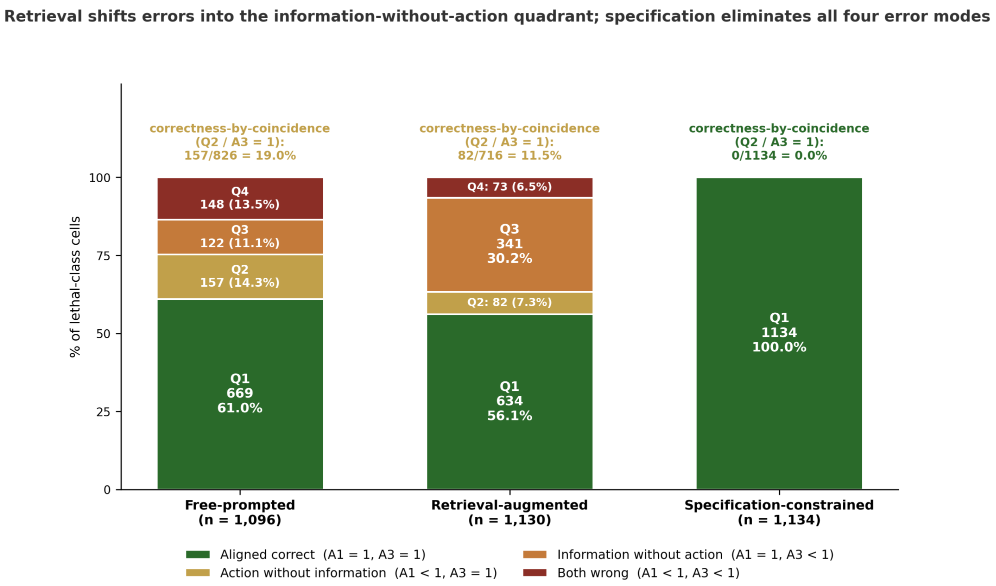


Lethal-class subset of the locked 3-of-3 replicate three-arm dataset. Quadrants defined on per-cell A1 (phenotype) and A3 (clinical safety action) scores. Annotation above each bar: correctness-by-coincidence rate (Q2 over A3 = 1 subset).

**Figure S4. Correctness by coincidence**. Per-condition decomposition of lethal-class cells into the four A1 (phenotype) by A3 (safety action) quadrants. Correctness-by-coincidence, the right action reached through a wrong phenotype (Q2 as a fraction of all action-correct cells), is 19.0% free-prompted, 11.5% under retrieval and 0% under the answer-supplied control, which reaches the aligned-correct quadrant on every cell.
